# Resistance variation and bacterial interactions shape the adaptation of a genetically diverse bacterial population to antimicrobial treatment

**DOI:** 10.1101/2025.04.01.646401

**Authors:** Aditi Batra, Leif Tueffers, Kira Haas, Tabea Loeblein, João Botelho, Michael Habig, Daniel Schuetz, Gabija Sakalyte, Florian Buchholz, Ernesto Berríos-Caro, Hildegard Uecker, Daniel Unterweger, Hinrich Schulenburg

## Abstract

Bacterial infections are often polymicrobial and subject to the evolution of antimicrobial resistance (AMR). Existing knowledge on AMR in such polymicrobial infections usually relies on observational patient data, for which cause-effect relationships are difficult to infer, or on studying interactions between different bacterial species, ignoring the commonly encountered variation within species. Here, we therefore asked how mixed populations with strains from the same species evolve under antibiotic treatment. We used a genetically diverse population of the high-risk human pathogen *Pseudomonas aeruginosa,* and first identified strain variation in both AMR and pairwise bacteria-bacteria interactions, the latter ranging from beneficial, neutral, to competitive. Using experimental evolution, we subsequently demonstrate that the response to selection by different antibiotic treatments is significantly influenced by AMR strain variation, bacterial interactions, and also spatial population structure. Moreover, *de novo* AMR evolution was additionally impacted by variation in resistance rates towards the two considered antibiotics. A second evolution experiment emphasized the central role of strain variation and bacterial interactions in determining the evolutionary outcome. We conclude that ecological dynamics in genetically diverse pathogen populations are key for our general understanding of infection characteristics and AMR evolution, and, therefore, deserve particular attention during treatment of polymicrobial infections.

## INTRODUCTION

Antimicrobial resistance (AMR) is an eco-evolutionary problem at its core^1^. Bacterial adaptation in response to antibiotic usage has resulted in widespread AMR, rendering many drugs ineffective^2^. As a consequence, we find ourselves in the middle of an AMR crisis, with approximately 4.95 million deaths worldwide associated with AMR in 2019^3,4^.

The key to countering AMR evolution is to understand how it happens. AMR evolution is usually a consequence of antibiotic therapy^5^ and can emerge within a few days of treatment^6–8^. AMR may be shaped by the number of infecting strains and species and thus be driven by a combination of ecological and evolutionary processes. Single strains dominate some infections, such as those related to sepsis^9^ or uncomplicated urinary tract infections^10^. In these infections, available resistance and pathogen evolvability shape AMR. Many other diseases can be polymicrobial. Infections in the abdominal cavity such as acute cholangitis^11^ or acute pancreatitis^12^ are frequently caused by subsets of the gut microbiome. The lungs of cystic fibrosis (CF) patients are often infected by multiple strains of *Pseudomonas aeruginosa* (PA)^13–16^. Based on this background, we here ask how does AMR evolve in such polymicrobial infections?

To date, almost all work on AMR in polymicrobial infections is based on observational data collected from patients^15,17,18^, for which the inference of cause-effect relationships is often a challenge. The few experimental studies on the topic focused on interactions between different bacterial species^19–22^, thus neglecting the variation commonly encountered within single species^13–16^. Such intra-specific genetic variation may increase the likelihood of pre-existing AMR^23^ or promote AMR spread through HGT^13^. Intra-specific variation may further associate with beneficial, neutral, or competitive interactions between strains^24^, which in turn can affect the spread of AMR, contingent on the exact nature of the interaction^25,26^. Spatial structure is an additional factor, particularly relevant for lung infections^27^, that can lead to independently evolving sub-populations^28^, thereby facilitating AMR emergence and spread^29^. Overall, intra-specific genetic variation is likely a critical factor in shaping AMR in infecting microbe populations, yet, to date, its exact role on AMR evolution is largely unexplored.

To address these current knowledge gaps, our study used an experimental approach to assess how AMR emerges and spreads in a genetically diverse pathogen population and to what extent pathogen evolution is influenced by standing genetic variation in AMR, microbial interactions, HGT, and spatial structure. Our genetically diverse population (gdPop) contained a mixture of 12 strains of the human opportunistic pathogen PA as a model. We first characterized variation in AMR and microbial interactions within the gdPop. Thereafter, the gdPop was subjected to an evolution experiment with different antibiotic treatments and two different transfer protocols, simulating two levels of spatial structuring. We analysed growth dynamics and strain diversity during experimental evolution, and characterized evolved AMR, genome sequence changes, and HGT at the end of the experiment. The insights from these measurements were tested and validated with an additional independently performed evolution experiment and also fluctuation assays to infer rates of AMR emergence.

## RESULTS

### gdPop strains vary in their antibiotic resistance, pairwise interactions, and HGT potential

The gdPop consisted of 12 strains from the previously characterized mPact strain panel^30,31^, covering the known genomic diversity of PA (Fig. 1A). Before performing the evolution experiment with the gdPop, we experimentally characterized the strains’ antibiotic resistance profiles and their pairwise interactions with one another. We found that the strains differed in their resistances to gentamicin (GEN) and piperacillin/tazobactam (PTZ), the two antibiotics used in our evolution experiments (Fig. 1B, Fig. S1). Pairwise bacterial interactions between the strains were quantified in the absence of antibiotics as growth in conditioned media (i.e., the supernatant of a culture of one of the bacteria). The reduced growth of the recipient strain A in donor strain B’s conditioned medium indicates a negative interaction while increased growth indicates a positive one. We found that negative interactions dominated the interaction matrix (Fig. 1C). Of the 144 one-way interactions quantified, 69% reduced growth of the recipient strain, 8% had no effect, and 22% increased recipient growth. We further calculated two types of cumulative interaction effects, focusing either on the cumulative effect of a focal donor strain on all others (focal −> others) or vice versa (others −> focal). These cumulative measures revealed substantial variation across the gdPop (Fig. 1D). As a last point, we note a high potential for HGT for the gdPop strains, demonstrated by our previous work that identified numerous integrative conjugative elements (ICEs), plasmids, as well as other mobile genetic elements for the strains^30^.

**Figure 1.**
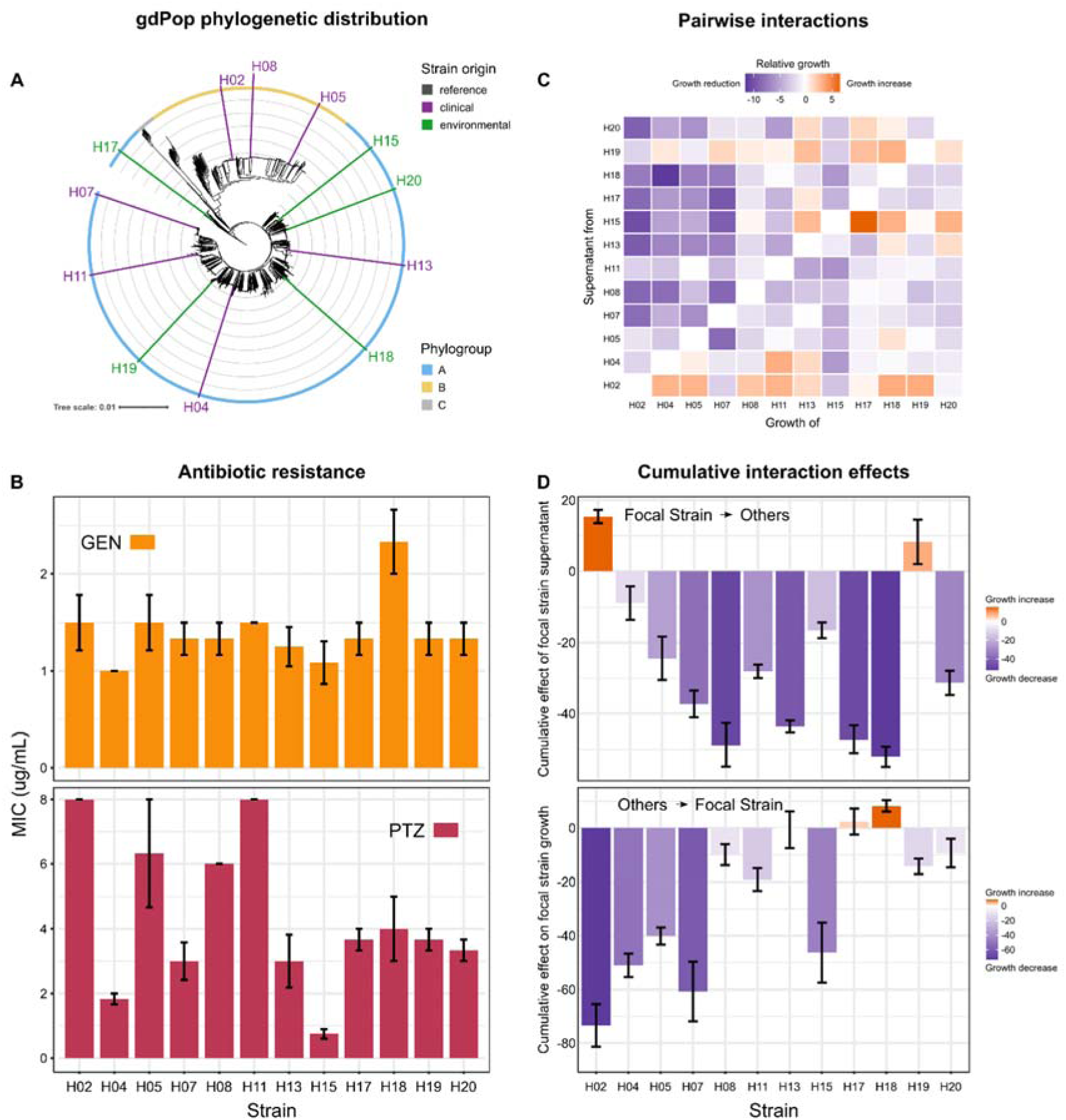
Strain characteristics of the *P. aeruginosa* genetically diverse population (gdPop). **(A)** The gdPop is composed of 12 strains from the major *P. aeruginosa* clone type (mPact) strain panel. These strains belong to the two major phylogroups of the species, cover a substantial part of the species’ genomic diversity, and represent both clinical and environmental isolates. **(B)** Resistance of the gdPop strains to gentamicin (GEN, in orange) and piperacillin/tazobactam (PTZ, in deep red). Variation in resistance among strains is shown as minimum inhibitory concentration (MIC) and found to be stronger on PTZ than GEN. Bars show the mean and error bars the standard error of the mean (SEM, n=3). **(C)** Pairwise interaction matrix of the 12 gdPop strains. A negative interaction was defined as reduced growth of recipient strain A in donor strain B’s conditioned medium (blue color) while increased growth indicated a positive interaction (red color). Relative growth was calculated by dividing growth in non-self conditioned medium with growth in self-conditioned medium (n=3). **(D)** The interaction matrix was used to calculate the effect of each individual focal strain’s supernatant (donor strain) on the growth of other strains (recipient strains) and vice-versa. The summation of the effect of focal strain supernatant on all other strains (one row in the matrix in C) gave the cumulative effect of focal strain supernatant (focal −> others). The summation of the effect of other supernatants on focal strain growth (one column in the matrix) gave the cumulative effect on focal strain growth (others −> focal). Both parameters showed comprehensive variation among strains. Bars show the mean and error bars the SEM (n=3). Data provided in supplementary data tables 1 and 2.

### Population mixing enhances adaptation under antibiotic selection

We evolved the gdPop under various antibiotic selection conditions to test the impact of standing genetic variation, bacterial interactions, and HGT on resistance evolution. Six treatments were included – two monotherapies, two switching treatments, a combination treatment, and a no-drug control (Fig. 2B). GEN and PTZ were chosen because they show drug synergy in combination^32^, reciprocal collateral sensitivity^33^, and directional negative hysteresis from PTZ to GEN (unpublished data) in PA (Fig. 2A) – all factors which should improve the efficacy of switching and combination treatments^32,34,35^. To assess the importance of spatial structure, we used two separate transfer protocols, resulting in two meta-treatments (Fig. 2C). The no-mixing meta-treatment involved a well-to-well correspondence between old and new plates at each transfer, thereby simulating spatially separated populations. In the mixing meta-treatment, all bacteria from replicate wells of a particular treatment were mixed in equal proportions at the end of each growth season, followed by transfer of this mixture to the fresh plates, thereby simulating the lack of spatial structuring. The experiment included 15 transfers with the last day in drug-free media to assess population survival (Fig. 2B). The dynamics of evolutionary adaptation were quantified during the evolution experiment using two parameters: (i) Time to adaptation was the time taken until a population under antibiotics had reached the same biomass (measured as optical density, OD) as the no drug control, while (ii) the robustness of adaptation was calculated as the number of growth seasons the population then remained at biomass levels of the no-drug control.

**Figure 2.**
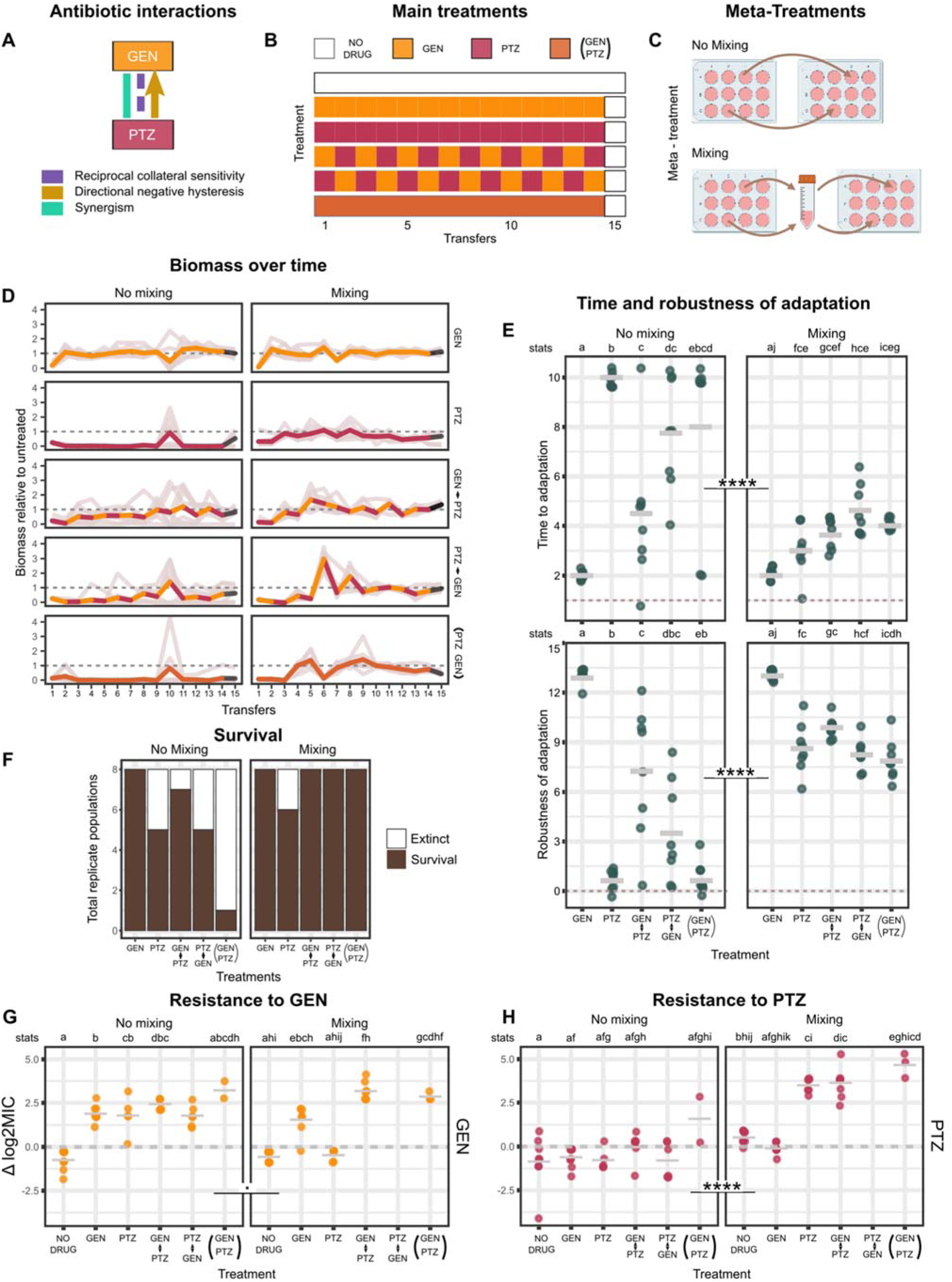
Experimental evolution of the gdPop under antibiotic selection. **(A)** Interaction among the two antibiotics used. Gentamicin (GEN) and piperacillin/tazobactam (PTZ) are known from previous work to interact synergistically towards *P. aeruginosa* (green color)^32^, to produce reciprocal collateral sensitivity to one another upon evolution of resistance to each one of the antibiotics in *P. aeruginosa* (purple color)^33^, and PTZ exposure sensitizes *P. aeruginosa* cells to treatment with GEN (i.e., negative hysteresis; yellow color; unpublished work; ^34^). Based on these characteristics, these antibiotics impose high selective constraints on evolving *P. aeruginosa* populations. **(B, C)** To test the impact of standing genetic variation, bacterial interactions, and horizontal gene transfer (HGT), the gdPop was evolved under six different specific treatments (B), including monotherapies, switching treatments (i.e., alternation of GEN and PTZ, starting with either of the drugs in separate treatments), a combination treatment, and a no-drug control. To assess a potential influence of HGT, an additional meta-treatment **(C)** was imposed on all six specific treatments (shown in **B**). The no-mixing meta-treatment involved a well-to-well correspondence during transfer after each growth season. The mixing meta-treatment involved mixing of all replicate populations within a particular specific treatment (shown in **B**) and inoculation of the next plate from this mixture. The latter meta-treatment served to limit the potential loss of strains due to genetic drift and to encourage HGT. Each individual treatment was run in 8 replicates. **(D)** Growth dynamics during the evolution experiment. Biomass (OD_600_) relative to the no-drug control is plotted. Light gray lines are the replicate populations, and the thicker, coloured line is the mean. Colours correspond to the specific treatments (shown in **B**). n=8 per treatment. The dotted line indicates growth levels for the no-drug control. **(E)** Adaptation response of the gdPop across specific treatments and meta-treatments. Adaptation was quantified using two parameters, the time to adaptation (i.e., the number of transfers until the growth level of the no-drug control was reached), and robustness of adaptation (i.e., the number of transfers the population reached the growth level of the no-drug control after having adapted once). Each data point is one replicate. Bar is the mean of all replicates per specific treatment. **(F)** Number of surviving populations per specific treatment out of a total of eight. **(G, H)** Log_2_MIC fold change of evolved gdPop relative to the ancestral gdPop for PTZ and GEN. Frozen populations from season 14 were regrown and the MIC was measured using MIC test strips (Liofilchem). The number of recovered populations varied from 2-8 per treatment. No populations were recovered for the PTZ −> GEN treatment in the mixing meta-treatment. Dots are the individual replicate populations while the bar is the mean. The dotted line represents the ancestral log2MIC. The statistical analyses of the adaptive responses in **E, G, H** were based on a permutation ANOVA, followed by Pairwise Wilcoxon Rank Sum posthoc tests to assess differences among specific treatments. Statistically significant differences between specific treatments are indicated by different letters, given above each of the panels (see line denoted “stats”). Overall, mixing increased the adaptive response, while reduced responses were mainly observed for treatments involving PTZ. Detailed results on the statistical analysis of the data is given in Tables S1-S9. All data for the figure panels is provided in supplementary data tables 3-7.

The inferred evolutionary dynamics varied significantly between the two meta-treatments and the individual treatments (Figs. 2D, 2E, randomized ANOVA, *p*<0.0001 for both parameters). Under no-mixing conditions, adaptation to GEN monotherapy happened in 2 days, while PTZ monotherapy exerted stronger selection with bacteria unable to adapt until the end (Fig. 2D). The switching treatments increased the time to adaptation compared to the GEN monotherapy but not to the PTZ monotherapy. The combination treatment also delayed adaptation. Populations that took longer to adapt also appeared to have a less robust adaptation. Adaptation was faster and more robust under the mixing conditions. Most strikingly, the gdPop adapted significantly faster to PTZ monotherapy and the combination in the mixing rather than the no-mixing meta-treatment (Fig. 2D). The mixing meta-treatment was also associated with a significant increase of population survival (logistic regression, *p*=0.001; Fig. 2F).

We also characterized AMR of the evolved populations using MIC test strips (Fig. 2G and H) and broth microdilution (Fig. S1B). PTZ resistance under no-mixing conditions was similar and not higher than that of the ancestor (Fig. 2H). In the mixing meta-treatment, the no-drug treatment and GEN monotherapy remained at ancestral PTZ MIC level while the others were significantly higher. Resistance to GEN was observed in all antibiotic treatments in both mixing and no-mixing meta-treatments except for PTZ monotherapy under mixing. Overall, mixing meta-treatment increased survival, adaptation, and AMR. Evolutionary adaptation was observed to be most constrained when PTZ was included.

### Treatment type, standing genetic variation, and bacterial interactions determine strain abundance under antibiotic selection

We next assessed to what extent the treatments and meta-treatments affected strain composition of the evolving gdPop. We used strain-specific PCRs on three randomly chosen replicate populations from transfers 4, 7, 11, and 13 (Fig. S2) and calculated a Shannon diversity index for each treatment combination. We identified temporal variation in Shannon diversity within treatments (Fig. 3A), and a significantly higher Shannon diversity under mixing than no-mixing conditions (Fig. 3C, GLMM, p <0.001).

**Figure 3.**
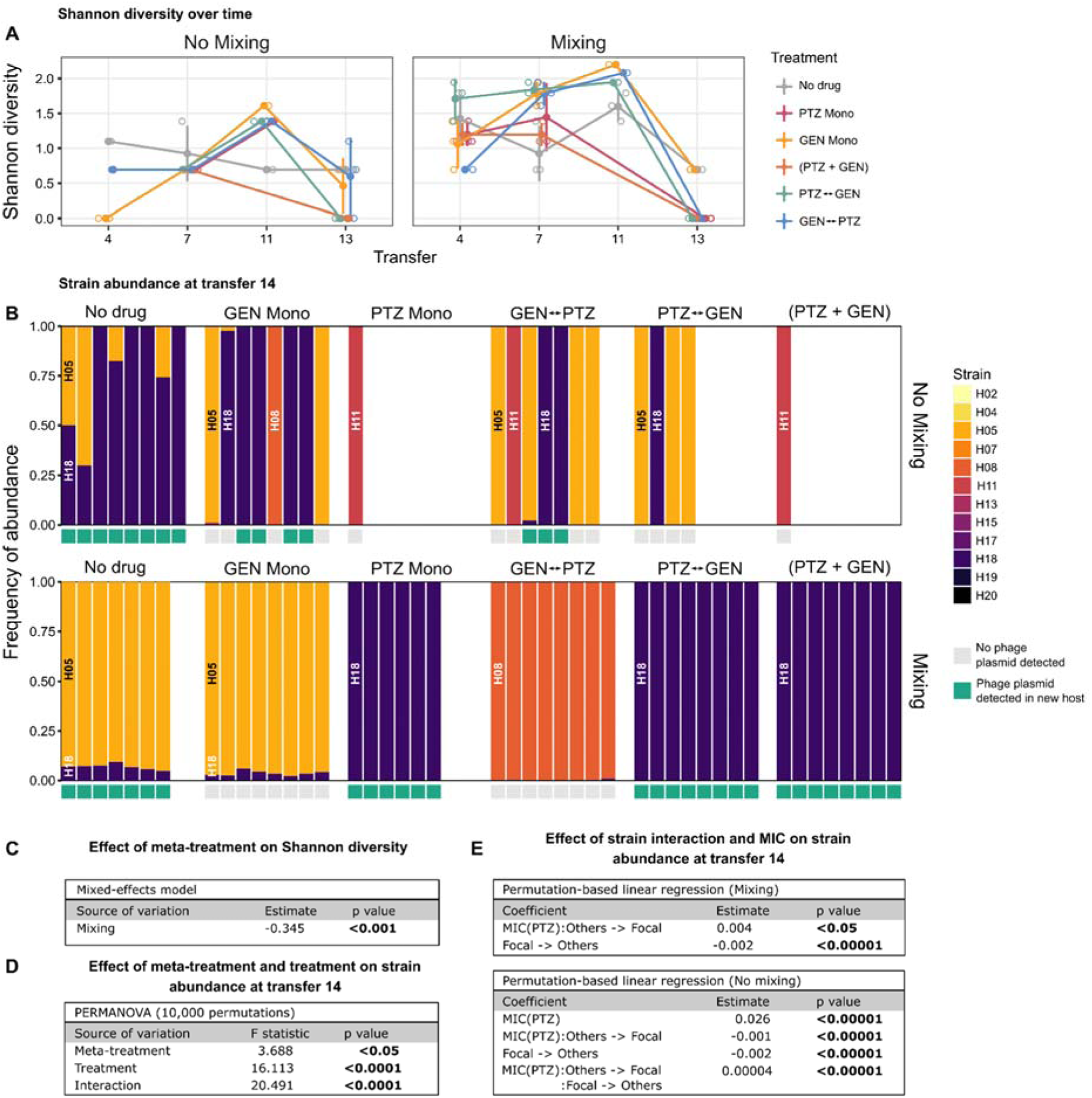
Strain diversity and horizontal gene transfer (HGT) in the gdPop. **(A)** Strain diversity during experimental evolution. Strain-specific PCRs (Table S10) were performed on three replicate populations from transfers 4, 7, 11, and 13 to obtain presence/absence data for the 12 strains, followed by calculation of Shannon diversity indices for each treatment and time point. Open circles are the individual replicates, and filled circles represent the mean. Error bars show standard deviation. **(B)** Whole genome sequences for populations from transfer 14 were used to determine strain abundance. Each bar corresponds to a replicate population. Strains are indicated by the different colours (see left). Empty columns refer to populations that went extinct (cf. Fig. 2F) or could not be recovered for genome analysis. HGT was only identified for a phage plasmid and indicated by green boxes right below the individual bars. **(C)** Results of the generalized linear mixed model used to assess the impact of meta-treatment on diversity. Mixing significantly increased strain diversity in treatments. **(D)** Results of the PERMANOVA used to test the effect of experimental design on strain abundance at transfer 14. Meta-treatment, treatment, and their interaction had a significant effect. **(E)** Results of the permutation-based linear regression used to test the effect of pre-existing resistance and bacterial interaction parameters on strain abundance, assessed for the two meta-treatments separately. Only significant coefficients are shown, indicating that both types (i.e., pre-existing resistance and interaction) significantly impacted abundance. Detailed results of statistical analyses are given in Tables S11-S14. The data is provided in supplementary data tables 8 and 9.

We further assessed strain abundance and HGT using whole genome sequence data for populations from transfer 14 (Fig. 3B, Fig. S3). Only 4 out of initially 12 strains were still present at the end, and usually one strain dominated a particular population. Strain abundance significantly varied with the meta-treatment, treatment, and their interaction (Fig. 3D, PERMANOVA, *p*<0.05 for all). We next tested if strain abundance was associated with starting strain MIC and bacterial interaction parameters, as determined above prior to the evolution experiment (Fig. 1B-1D). We found that the cumulative effect of focal strain conditioned media on others (focal −> others) significantly decreased strain abundance (Fig. 3E, PERMLM, *p* < 0.0001) for both mixing and no-mixing meta-treatments. This is consistent with the observation that most strains negatively affected others. The MIC on PTZ increased strain abundance under no-mixing conditions (PERMLM, *p*<0.0001). The interaction between PTZ MIC and the cumulative effect on focal strain growth (others −> focal) also significantly affected abundance in both meta-treatments (Fig. 3E, PERMLM, *p*<0.0001). Finally, the three-way interaction between the PTZ MIC, focal −> others, and others −> focal was a significant predictor of strain abundance in the no-mixing meta-treatment (Fig. 3E, PERMLM, *p*<0.0001).

If we consider the four predominant strains, then these observed patterns can be explained most easily for H18, which receives the strongest benefit from the interaction with other strains (Fig. 1D, bottom panel), while also outcompeting the others (Fig. 1D, top panel). Another strain, H08, mainly dominates in the GEN->PTZ switching treatment under mixing conditions. This is possibly because H08 benefits from many of the pairwise interactions and is one of the stronger competitors (Fig. 1C, 1D). Strain H11 dominates in single replicates in treatments with PTZ under no-mixing conditions (Fig. 3B). This strain benefits from some of the interactions (Fig. 1C, 1D, bottom panel) and, importantly, shows one of the highest resistance levels towards PTZ (Fig. 1B, bottom panel). Finally, the strain H05 dominates the no-drug control and the GEN monotherapy treatments under mixing conditions and also a few replicate populations of different multi-drug treatments under no-mixing conditions (Fig. 3B). This pattern cannot be easily explained with the available data, as this strain does not perform very well in the pairwise interactions and it also only shows somewhat higher resistance levels towards PTZ, but not GEN, when compared to the other strains (Figs. 1B-1D). Overall, we conclude that strain abundance during antibiotic selection was influenced by the type of treatment, the pre-existing resistance to the antibiotics as well as the interactions between bacteria.

We further identified HGT, in all cases of a phage plasmid from strain H13, which spread to new hosts in 59% of the populations (Fig. 3B, Fig. S3). The genes located on this phage plasmid have not been reported to affect antibiotic resistance (Fig. S3). Therefore, the spread of the phage plasmid is likely a consequence of selfish behavior and unlikely to have contributed to AMR evolution.

### Increases in AMR are associated with mutations in known AMR genes and variation in rates of *de novo* resistance emergence

We noted increases in GEN and PTZ resistance in populations dominated by the same PA strain. For example, the strain H05 dominated in the control treatment and the GEN monotherapy under mixing conditions, whereby we only observed an increased GEN resistance in the latter but not the former (Figs. 2G, 3B). Similarly, H18 was most abundant in PTZ monotherapy under mixing conditions, but also prevailed in several replicates of the control treatment under no-mixing conditions; an increased resistance was only recorded for the former but not the latter (Figs. 2H, 3B). These observations suggest that the resistance increase is not only due to changes in gdPop strain composition but additionally mediated by newly emerged resistance mutations. To assess this point, we characterized the distribution of mutations in known PA AMR genes across treatments and meta-treatments. We found that AMR gene mutations only occurred in treatments with antibiotics and, in this case, in almost all of the replicate populations, whereas no AMR gene mutation was identified for the no-drug controls (Fig. 4A). Interestingly, the mixing meta-treatment led to the uniform spread of the same variants across replicate populations of a particular treatment, including variants in the gene *ampR* whenever a treatment contained PTZ and also variants in *pmrB* or *parS* for treatments with GEN (Fig. 4A). The no-mixing meta-treatment produced more variation in the favored AMR gene mutations, including variation in the detected mutations in *pmrB* (Fig. 4A). Based on these results, we conclude that the mixing meta-treatment favored a uniform selection of resistance variants, most likely those with highest competitive fitness, whereas the observed variation in selected variants under no-mixing conditions was likely influenced by chance effects.

**Figure 4.**
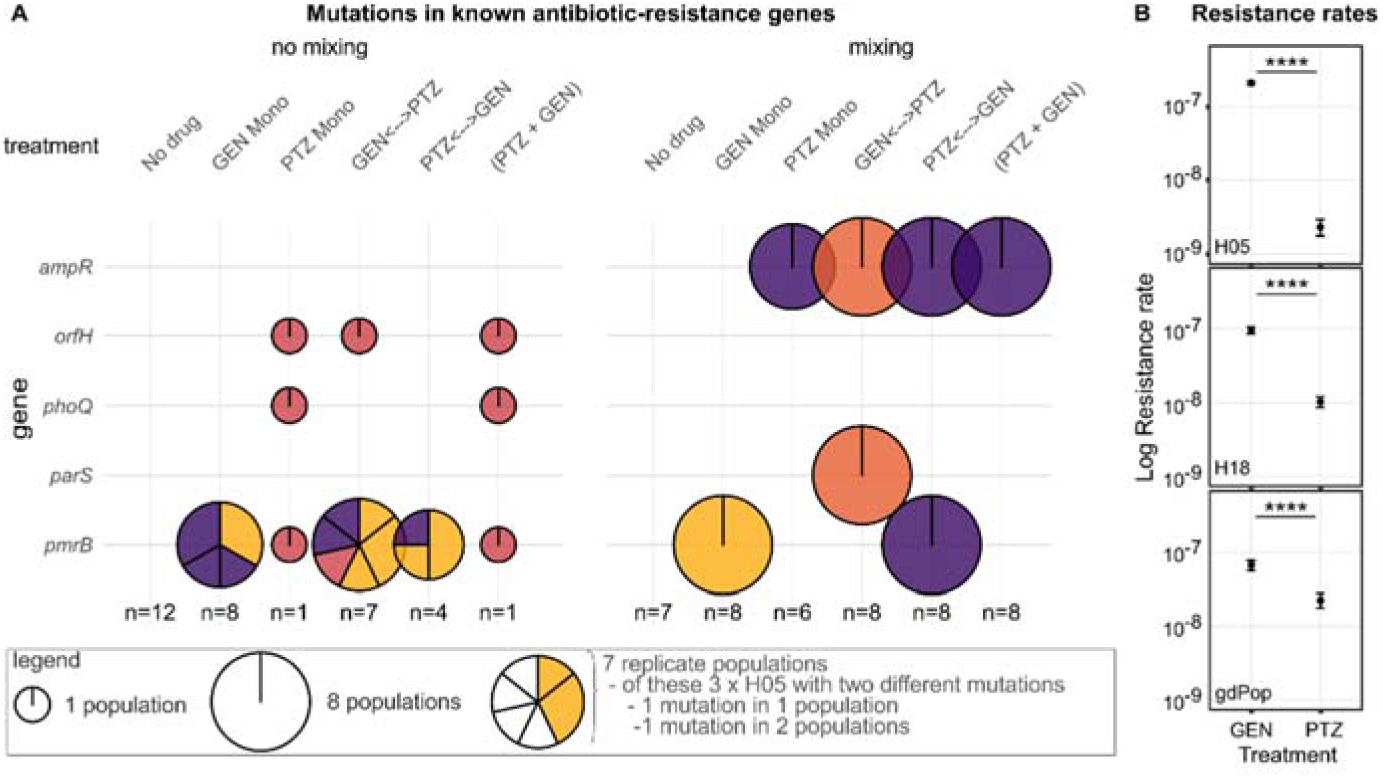
The distribution of mutations in known resistance genes and variation in resistance rates towards GEN and PTZ. **(A)** Mutations in known AMR genes identified for populations from different treatments at the end of the evolution experimental. The known AMR genes are given along the vertical axis, while the different treatments and meta-treatments are shown horizontally. The size of the circles corresponds to the number of replicate populations with a variant in a particular gene. Within each circle, different background genotypes are indicated by color, following the same color scheme as used in Fig. 3B (H05, yellow; H08, light red; H11, dark red; H18, purple). Different mutations within a specific genotype are shown as separate pieces. All no-drug control samples as well as two replicate populations of GEN monotherapy under no-mixing conditions did not show a mutation in a known AMR gene, while all other populations did. For the no-mixing treatment, the single populations from PTZ monotherapy and combination treatment as well as one of the seven populations from switching treatments starting with GEN simultaneously had mutations in at least two genes. For the mixing meta-treatment, all replicate populations from the switching treatments had mutations in exactly two genes. All remaining replicate samples showed a single mutation in a known AMR gene. **(B)** Resistance rates towards GEN or PTZ. Resistance rates were inferred using the fluctuation assay with either the single strains H05 or H18 or the whole gdPop. The results are shown as the mean of two independent runs. Error bars represent confidence intervals (CI95). The statistical comparison between antibiotics was based on Likelihood Ratio Tests. The four stars indicate a significantly lower resistance rate towards PTZ than GEN (*p*<0.0001; detailed statistical results in Table S15). The data for this figure is provided in supplementary data tables 10-12.

Although these findings can explain the observed resistance increases, they cannot explain the lack of resistance evolution against PTZ (Fig. 2G) or the increased extinction rates in treatments with PTZ (Fig. 2F). We postulated that these results may be due to a small mutational target size for PTZ resistance and thus a small rate of spontaneous PTZ resistance mutations. We tested this idea with standard fluctuation assays^36,37^ for H05, H18 (the two dominant strains at the end of evolution), and gdPop on both GEN and PTZ. We found that the resistance rates towards PTZ were indeed significantly lower than those towards GEN – consistently in both tested strains and the gdPop (Fig. 4B, Likelihood Ratio tests, *p*<0.0001 for all). Therefore, the comparatively lower PTZ resistance rates appear to underlie the observed lower rates of resistance evolution and the higher extinction rates during experimental evolution in PTZ-containing treatments.

### Strain diversity increases population survival under antibiotic selection in a second independent evolution experiment

Since the mixing meta-treatment maintained a higher strain diversity across time (Figs. 3A, 3C), the observed higher rate of adaptation could have been due to (i) a reduced random loss of strains, including those that already express resistance or show a high potential to evolve resistance, and/or (ii) the maintenance of beneficial bacterial interactions that somehow facilitate resistance. The mixing conditions could have further enhanced adaptation by producing an overall larger population size (1 well in no-mixing vs 8 connected wells in mixing) and thus a higher likelihood for the occurrence of favorable new mutations. To further explore the relevance of these alternatives, we performed a second independent evolution experiment. To test the effect of diversity, the evolution experiment was set up with either only single strains (H05 or H18) or the gdPop. The two considered single strains were most frequently observed at the end of the first evolution experiment, indicating their genetic potential to be maintained during experimental selection. If these were unable to adapt, it would suggest that adaptation in the first experiment was not due to the retention of successful strains, but rather resulted from selective benefits arising from bacterial interactions. To explore a possible effect of population size, the experiment was carried out in two volumes, two and four milliliters, whereby we ensured that antibiotic selective pressure (i.e., antibiotic concentrations) and also cell density was identical across the two volumes (Fig. S4). The bacterial populations were then subjected to three selective treatments across five days (Fig. 5A; 20 replicates per treatment combination). During the evolution experiment, we monitored bacterial growth by measuring biomass (OD_600_) (Figs. 5B, 5E) and colony-forming units (CFU/ml) (Fig. S5).

**Figure 5.**
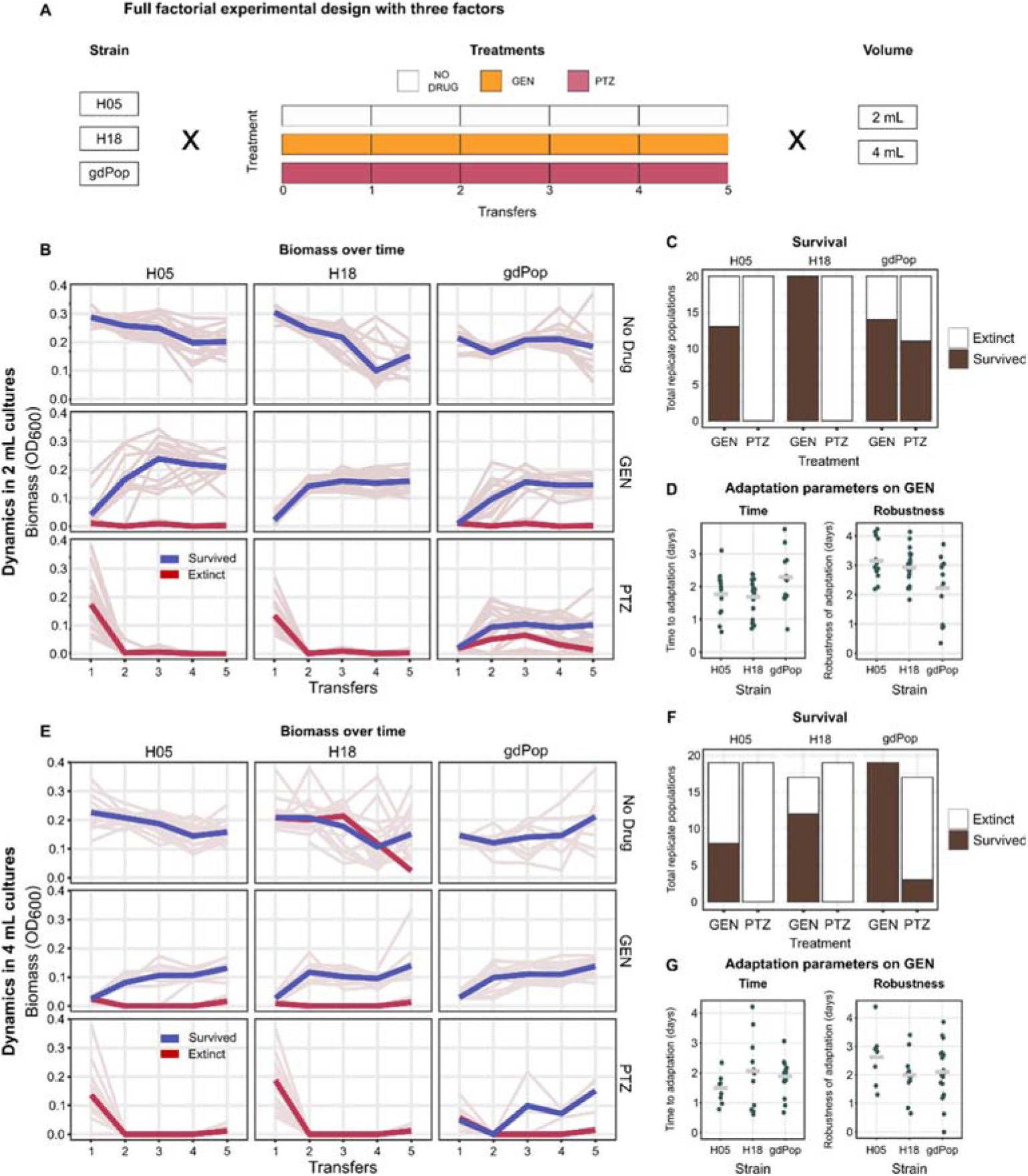
Validation evolution experiment with gdPop and two single strains in different volumes under antibiotic selection. **(A)** Design of the repeat evolution experiment to explore a possible influence of population size. Based on the limitations of comparable growth plates, population size was varied by evolving bacteria in two different volumes, 2 ml and 4 ml. This set-up allowed us to double the population size in the larger volume, while ensuring the same density of bacterial cells across the two volumes. This setup was used for two single gdPop strains, H05 and H18 (the two most frequent survivors of the main evolution experiment), and the whole gdPop. Bacteria were subjected to three treatments, two antibiotic monotherapies, and a no-drug control, over five days. Each treatment combination was replicated 20 times. **(B, E)** Growth dynamics during the evolution experiments for either small (B) or large (E) volumes. Biomass (OD600) over time is plotted. Light gray lines are the replicate populations, and the thicker, coloured lines are the means of the surviving (blue) and extinct (red) replicates. n =17-20 per strain, treatment, and volume combination. **(C, F)** Surviving populations per strain and treatment for small (C) or large (F) volumes. The total height of the bar represents the total replicates tested (with a possible maximum of 20). Black and white represent surviving and extinct replicate populations, respectively. Volume, strain, and antibiotic were all found to significantly affect survival (logistic regression). The higher volume and PTZ negatively affected survival (*p*<0.05 and *p*<0.0001, respectively) while strain diversity (gdPop) significantly increased survival (*p*<0.0001) **(D, G)** Adaptation parameters for populations growing in the GEN treatment for either small (D) or large (G) volumes. Adaptation was quantified as the time to adaptation and robustness of adaptation (see legend to Fig. 2). Each data point is one replicate. Bar is the mean of all replicates. The time taken to adapt to GEN only showed a trend for the interaction term of Volume and Strain (Permutation ANOVA, *p*=0.0565). Robustness of adaptation was significantly affected by both Volume and Strain (Permutation ANOVA, *p*< 0.01 and *p*<0.05 respectively), with a higher volume resulting in less robust adaptation and H05 having significantly more robust adaptation than gdPop (Wilcoxon Sum Rank Tests, *p*<0.05). Detailed results for the statistical analysis is provided in Tables S16-S19. All data for the panels of this figure is provided in supplementary data tables 13-21.

We found that strain diversity but not strain identity or volume influenced the dynamics of adaptation to antibiotics. Under GEN monotherapy, the proportion of extinct replicate populations varied between strains (Figs. 5C, 5F) and further depended on the initial CFU/mL that usually show some variation under the used experimental conditions. In this case, we now observed that replicates with low initial CFU/mL were more likely to go extinct (Fig. S5). On PTZ, strain diversity positively impacted survival (logistic regression, *p*<0.05): all single-strain replicates went extinct, whereas at least some gdPop replicates survived (Fig. 5C, 5F). We next assessed the pattern of adaptation for the antibiotic treatment with little extinction, the GEN treatment, using the parameters of time to adaptation and the robustness of adaptation (Figs. 5D, 5G). The time to adaptation was not significantly different between volumes, but it showed a statistical trend of a difference among the three strains and the interaction of strain and volume (randomization ANOVA, *p*=0.59 for volume, *p*=0.08 for strain and *p*=0.056). The robustness of adaptation was significantly different between volumes and the strains (randomized ANOVA, *p*<0.05 for both).

Taken together, our combined results strongly suggest that the higher survival and faster adaptation observed in the mixing meta-treatment was a consequence of higher strain diversity. The increased diversity enhanced adaptation of the bacterial populations not by limiting the random loss of strains with the genetic potential to survive, but most likely by retaining bacterial interactions that contribute to AMR.

## DISCUSSION

Many infections are polymicrobial and often include strain variation within species^11–13,15^. In our study, we used a controlled experimental approach to demonstrate that strain diversity increases population survival under antibiotic selection (Figs. 4D, 4G), most likely as a consequence of strain interactions within the population. Indeed, we found that the selectively favoured strains at the end of evolution are those that benefit the most from interacting with other bacteria and have high pre-existing resistance (Fig. 6). In the following, we will discuss the importance of spatial structure, bacterial interactions, and standing genetic variation for our findings.

**Figure 6.**
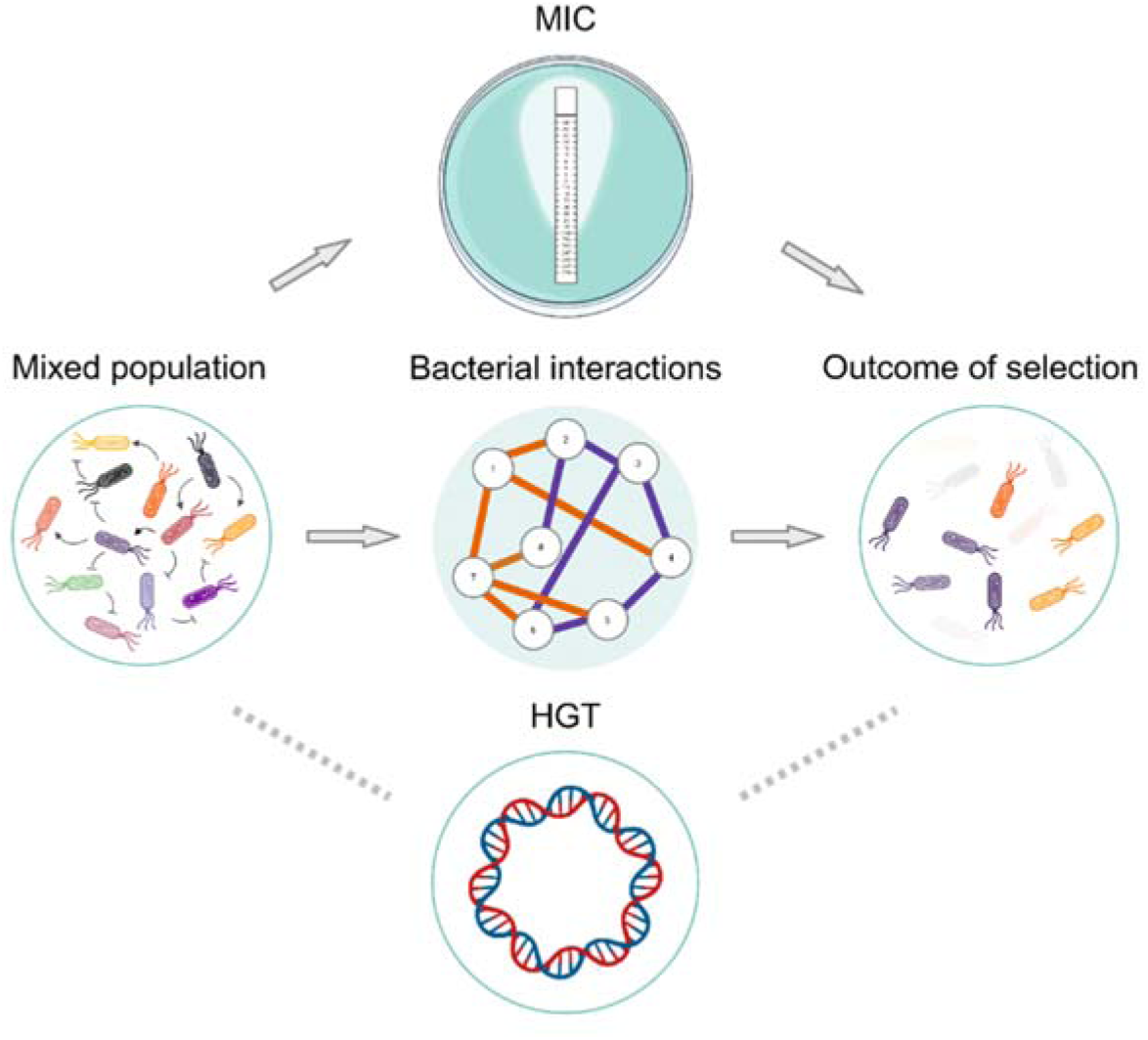
Bacterial interactions and minimum inhibitory concentration (MIC) determine selection outcome. Summary of the main findings of our study. We assessed to what extent variation in antibiotic resistance (measured as the minimum inhibitory concentration, MIC), bacterial interactions, and horizontal gene transfer (HGT) shape adaptation of a genetically diverse population of *Pseudomonas aeruginosa* to antibiotic selection. We found that both pairwise bacterial interactions and the MIC are key predictors of strain composition. Bacterial interactions were further found to contribute to population survival through their beneficial effect on the focal strain H18.

Spatial structure plays a key role in the adaptation of microbial communities, affecting biodiversity and competition^38^. The influence of spatial structure on adaptation has been addressed comprehensively using theoretical approaches^39–41^. In structured settings, populations are separated and beneficial mutations spread independently across subpopulations, potentially extending the duration of selective sweeps^42^, and facilitating the coexistence of multiple genotypes^43,44^. Spatial structure has indeed been shown to favour genetic diversity in subpopulations under migration and serial dilution^28^. However, spatial structuring can also boost the spread of beneficial mutations when mutations are rare^45^. The no-mixing and mixing meta–treatments in our main evolution experiment represent a high and low level of spatial structure, respectively. Our findings show that, as expected, the more structured populations (i.e., no-mixing meta-treatment) show a decrease in the speed of adaptation (Figs. 2D, 2E), while also producing a larger diversity of sequence variants between populations (Fig. 4A left). At the same time, the less structured conditions (i.e., mixing meta-treatment) lead to a uniform spread of sequence variants across populations (Fig. 4A right) and an at least initial increase in strain diversity (Fig. 3A), the latter possibly because of a reduced likelihood of random strain loss.

The higher strain diversity under mixing conditions is associated with higher adaptation rates (Figs. 2D, 2E, 3A, and especially 5B-G), possibly due to maintaining beneficial interactions between the bacteria. Bacterial interactions were previously implicated in modulating antibiotic resistance^25,26^. For example, cross-feeding can decelerate *de novo* resistance evolution, when one partner always has to ‘wait’ for its cross-feeding partner to evolve higher resistance^19,46^. As another example, the presence of carbapenemase-producing *Stenotrophomonas maltophilia* allowed coexisting PA to survive and subsequently evolve imipenem resistance^20^. Studies assessing the impact of a more complex community on resistance evolution are fewer and usually follow the dynamics of a focal strain within a community subjected to antibiotics. Interspecific competition increased the costs of resistance in focal *Escherichia coli* resulting in higher minimal selective concentrations^21^. However, a subsequent increase in antibiotic concentration led to the competitive release of the focal strain. Another study found that resistance evolution in focal *E. coli* was suppressed in human faecal communities^47^. To the best of our knowledge, our study is the first to demonstrate increased survival as a consequence of intra-specific bacterial interactions (Figs. 5C, 5F). Indeed, the ultimately dominant strain H18 (Fig. 3B) benefitted from the others the most while also outcompeting them (Fig. 1C, 1D). Thus, the overall distribution of interactions impacts how the whole population adapts to novel selective pressure, like that imposed by antimicrobials.

Strain abundance in our gdPop was significantly influenced by pairwise bacterial interactions as well as standing genetic variation in AMR (Fig. 3B, 3E). Our finding of the importance of microbial interactions is in line with previous work that showed that competition between pairs could predict survival in multi-species^48^ or multi-strain communities^49^. Moreover, standing genetic variation can also contribute to adaptation by providing the necessary diversity to adapt^50^. For example, in patients colonized by multiple PA strains, the pre-existence of resistant strains facilitated the spread of AMR^23^. Altogether, strain interactions as well as standing genetic diversity are important predictors of population dynamics.

The inability of bacteria to adapt to PTZ as compared to GEN was likely caused by a low rate of PTZ resistance mutations (Fig. 4B). The inclusion of this antibiotic in switching and combination treatments increased their efficacy (Figs. 2D-2F, 5B-5G). These findings are consistent with our previous demonstration that the inclusion of an antibiotic with low resistance rates in switching treatments with only beta-lactams increased bacterial eradication^51^. In the current study, we confirmed this effect, however, using different antibiotics than before and observed in both switching as well as combination treatments.

Predicting how populations or communities change over time is of particular interest in microbial ecology and additionally of high relevance in medical infectiology. Our work now demonstrates that the presence of strain diversity enhances the ability of bacteria to counter the high selection imposed by antibiotics. Thus, inter-strain interactions are as important as inter-species interactions in their effect on evolution. In turn, the consideration of these ecological processes is key to understanding antibiotic resistance spread in the widely occurring polymicrobial infections.

## Materials and Methods

### Strains

The genetically diverse population (gdPop) used for experiments consisted of 12 strains of *P. aeruginosa*. These were taken from the mPact strain panel representing the entire genomic diversity of the species^30,31^. Each of the 12 strains was grown separately in 10mL Luria Bertani (LB) medium for 20h at 37°C and 150 RPM following which they were mixed in a 1:1 ratio (v/v) to obtain the final mixed population. No statistically significant difference in the CFU/mL was observed at the end of 20h between the different strains (Kruskal Wallace Test, χ2=19, *p*=0.456, ref 28) This mixture was then used for experiments.

### Media and antibiotics

All experiments with the gdPop were conducted in either Luria Bertani (LB) or minimal M9 medium as broth or combined with agar (1.5%). M9 medium was supplemented with glucose (2 g/L), citrate (0.58 g/L), and casamino acids (1 g/L). Antibiotics were added when required. Antibiotics used included the aminoglycoside gentamicin (GEN, Carl Roth, Order No. HN09.1) and a combination of the beta-lactam piperacillin (PIP, Sigma Aldrich, Code: P8396-1G) and the beta-lactamase inhibitor tazobactam (TAZ, Sigma Aldrich, Code: T2820-10MG). Piperacillin and tazobactam (PTZ) were combined in an 8:1 ratio. Bacteria were incubated at 37°C.

### Experiments with conditioned media

To determine contact-independent pairwise interactions, we cultured the 12 strains separately in LB and M9 media overnight at 37 **°**C. After incubation, cultures grown in the M9 medium were filtered through 0.22 μm filters to generate bacteria-free supernatant. Strains grown in LB were sub-cultured until the mid-exponential phase and washed from the residual medium. We resuspended washed bacteria in the filtered M9 medium using the matrix-like format in a 96-well plate and normalized the initial OD to OD_600_ = 0.001. Cultures were incubated at 37 **°**C for 62 h with shaking (162 rpm). Measurements of OD_600_ were taken every 15 minutes with a microplate reader (Tecan, Spark). We performed 3 independent biological repetitions for each combination of strain and conditioned media. Area under the curve (AUC) measures were calculated using the R package *growthcurver.* Strain interactions were determined using the formula:

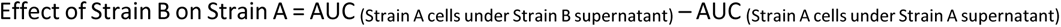

AUC values for the effect of Strain B supernatant on other strains’ growth (Focal strain −> Others) were summed up to obtain the cumulative effect of strain B’s supernatant. AUC values for the effect of other strains’ supernatant on Strain B’s growth (Others −> Focal strain) were summed to obtain the cumulative effect on Strain B’s growth.

### Minimum inhibitory concentration (MIC) measurement using MIC test strips

The MIC of both antibiotics for all strains (and later ancestral and evolved populations) was measured using the MIC test strips (Liofilchem)^52^. The individual strains were grown in 10 mL M9 for 18 h. Each strain was then diluted to an OD_600_ of 0.08 and spread onto a M9 plate using a cotton swab. MIC test strips (Liofilchem) were placed onto the plate and the plate incubated for 24h after which the MIC was read at the intersection of the zone of inhibition and the test strip. For testing the MIC of the gdPop, the 12 strains were grown individually in 10 mL LB for 20h and then mixed in equal proportions. This mixture was then diluted to an OD600 of 0.08 and spread onto a M9 plate using a cotton swab, followed by MIC inference as above using MIC test strips (Liofilchem). For PTZ, we observed bacterial growth in the shape of wings above the intersection of the zone of inhibition and the test strip (Fig. S1A). This is likely due to the paradoxical effect^53^. It is important to note that this effect was not observed on PTZ when individual strains were tested. The MIC was read where the top edge of the wings intersected the test strip. To measure MIC of the evolved populations from day 14, we thawed the frozen material from this timepoint, transferred 50 µL of the sample into 10 mL LB, followed by cultivation for 20 h at 37 °C and 150 rpm. After incubation, the cultures were diluted to an OD_600_ of 0.08 and then prepared for strip-based MIC inference as above. Change in MIC of the evolved population was calculated as MIC_evolved population_ - MIC_ancestor._

### OD-based dose-response curves

MIC test strips measure antimicrobial susceptibility based on growth on agar. To have a measure of antimicrobial susceptibility based on growth in liquid media, we carried out OD-based dose-response curves. For the ancestral population, the 12 strains were grown individually in 10 mL LB for 20 h after which they were mixed in equal proportions. 2 µL (2%) of this mixture was inoculated into the wells of a 96-well plate (100 µL total volume) containing a range of concentrations of antibiotics covering the MICs of the individual strains. The plate was incubated at 37 °C and 750 rpm for 24 h after which the OD_600_ was read in a BioTek plate reader. The area under the resulting dose-response curve was calculated using the *drc* package in R and was used as a measure of the antimicrobial susceptibility of the ancestral population. For measuring the antimicrobial susceptibility of the evolved populations, the populations frozen on day 14 were thawed and 50 µL was inoculated into 10 mL LB and then subjected to the same procedure described above. Change in antimicrobial susceptibility of the evolved population was calculated as antimicrobial susceptibility _evolved population_ - antimicrobial susceptibility _ancestor_.

### CFU-based dose-response curves

Dose-response curves (DRC) were carried out to determine the concentration of antibiotics reducing cell number by three orders of magnitude compared to the no drug control (Inhibitory concentration of 99.9% or IC99.9). Individual strains were grown overnight for 20 h in LB and then mixed in a 1:1 (v/v) ratio. 40 µL (2%) of this mixture was inoculated into the wells of 12 well plates containing a range of antibiotic concentrations chosen based on the MIC data of the individual strains. No drug controls and blanks were included. All treatments were randomized on the plate. Total volume in each well was 2000 µL. Plates were incubated at 37 °C and 900 rpm for 24 h. At the end of incubation, the plates were sonicated for 1 min to break any clumps of bacteria and these were spotted onto plates to determine CFU/mL. This was done for GEN, PTZ, and their combination (Fig. S1A). For the combination DRC, each concentration tested contained an equal mixture of the two antibiotics. CFU-based dose-response curves were repeated for the second evolution experiment in the new experimental settings required for the experiment.

### Strain-specific Polymerase Chain Reaction (PCR) and estimation of strain diversity over time

To detect which strains survived post-antibiotic treatment, strain specific PCRs were carried out. Each strain was uniquely identified by a primer pair (for a list of primer pairs see Table S1). Samples were taken at concentrations that reduced the cell number by three orders of magnitude and subjected to PCRs. PCRs included an initial denaturation of 45 seconds at 95 °C, followed by 30 cycles of annealing for 30 seconds at 55 °C and extension for 30 seconds at 72 °C, and then a final extension for 10 minutes at 72°C. To estimate strain diversity over time in the evolution experiment, strain-specific PCRs for each of the 12 strains were performed on 3 independent replicate populations from transfers 4, 7, 11, and 13. For the populations from transfer 13, DNA for the PCRs was isolated directly from the evolution experiment. For the rest, the frozen populations had to be regrown in LB for DNA isolation for the PCRs. Shannon diversity was calculated from the strain occurrence at these transfers for every combination of treatment and meta-treatment.

### Evolution experiment 1

The evolution experiment consisted of 6 treatments nested within 2 meta-treatments. The 6 treatments included 3 evolution-informed treatments (one combination and two sequential, each starting with a different antibiotic) and 3 controls (No drug, monotherapy with each antibiotic). Each treatment contained 8 replicate populations. Treatments were fully randomized on each plate. Each of the 12 strains were grown for 20 h in LB and then mixed in equal proportions (v/v). From independent mixtures bacteria were inoculated into 12 well antibiotic plates containing 2000 µL of medium. They were then grown for 24 h at 37 °C and 900 rpm, following which the plates were sonicated to break any bacterial clumps. 40 µL (2% v/v) was then transferred to another plate containing fresh medium. For the No-mixing meta-treatment, we ensured a 1-to-1 correspondence between wells during transfer. The ‘mixing’ meta-treatment was introduced to limit strain loss due to genetic drift and simulate spatial structuring. For this, all 8 replicates within a treatment were mixed together in one 50 mL tube and this mixture was used to inoculate all wells of this treatment for the next growth season. Each of the meta-treatments had 6 treatments resulting in a total of 12 treatments and 96 bacterial populations. Growth of the bacteria was followed by measuring endpoint OD_600_ over time. The experiment lasted a total of 15 days with 14 days of antibiotic exposure followed by 1 day of growth in drug free medium to assess population extinction. Populations were frozen in DMSO on days 4, 7, 11 and 14. Populations that grew after the removal of antibiotics were frozen on day 15.

### Calculation of population extinction and adaptation parameters

Both population extinction and adaptation of the surviving bacterial populations to the treatment was quantified at the end of the evolution experiment. A population was identified to be extinct if in the last growth season without drug its OD_600_ was less than 0.05. Two parameters were used for quantifying the pattern of adaptation. Time to adaptation is the time taken (in number of growth seasons) for the bacterial population under treatment to attain the same level of growth as those from the no-drug control. Robustness of adaptation is the number of growth seasons the bacterial population under treatment is able to maintain the growth level of the no-drug control (i.e., its OD_600_ was identified to be within the mean ± 2x standard deviation of the no-drug control in that growth season), after this level was first reached. This is a measure of how well adaptation was maintained once acquired.

### Determination of strain abundance, horizontal gene transfer, and sequence variants

DNA was isolated from the evolved populations at transfer 14 and used for Illumina HiSeq3000 short read sequencing. Quality control of the Illumina reads was performed using MultiQC v1.12^54^ with default parameters. To perform strain-level abundance estimation in our metagenomic samples using variation plots, we used StrainFLAIR v0.0.1 with default parameters^55^. Analysis of horizontal gene transfer was done using a subset of genomes sequenced with both short and long reads, using sequencing on an Illumina HiSeq3000 and PacBio Sequel II, respectively. Illumina short reads were assembled using SPAdes v3.15.5 in meta-assembly mode^56^. PacBio long reads were assembled using Flye v2.9.3-b1797 in metagenome assembly mode^57^. Short and long read assemblies were annotated using Bakta v1.9.2 with default parameters against the complete database^58^. We used the wild-type genomes from the gdPop collection and the short-read assemblies to build a pangenome using Panaroo v1.4.2^59^. We ran the analysis in strict mode, with the flag to remove invalid genes and a protein family sequence identity threshold of 90%. To investigate the genetic rearrangements of the phage-plasmid acquired from H13, we compared the relevant genomic regions in the wild-type genomes of H13 and H18 and the long-read assemblies of the metagenomic samples using clinker v0.0.20^60^ and a minimum alignment sequence identity of 90%. For samples with only short reads, the phage replicase from the phage-plasmid was compared using DIAMOND (version 2.1.9)^61^ with the blastx mode against all short-read assemblies.

The analysis of sequence variants was based on the short reads data. Reads were trimmed using Trim Galore (version 0.6.10) and then split between the twelve reference genomes using BBsplit (version 38.18), with ambiguous reads mapped to all reference genomes. Reads mapped to the dominant or non-dominant strains were aligned to their respective genomes using Bowtie2 (version 2.4.1)^62^, and SNPs were called using FreeBayes (version v1.3.2-dirty). SNPs and INDELs were filtered for a quality score >50 using BCFtools (version 1.14)^63^ and GATK (version 4.0.5.1). SNPs/INDELs detected in the ancestral strain, as well as those found when mapping reads from the non-dominant strain onto the reference strain, were removed using a combination of BCFtools and BEDTools (version v2.30.0). To account for the high proportion of non-dominant reads in samples NMDF1 and NMDF2 (both No Drug control), the quality threshold was increased to 150 upon visual inspection in these samples. The orthologues of known resistance genes (*mexY, fusA, mexN, csrA, nalC, ftsI, mexR, ampC, ampR, mpl, dacC, nalD, dacB, ompQ, galU, amrR, mexX, parR, parS, PA14_45870, phoQ, ampDh3, mraY, ampD, pmrB, cysQ, amgS, ampDh2, orfH*) involved in resistance against GEN or PTZ were identified using OrthoFinder^64^. SNPs and INDELs overlapping these genes were determined using BEDTools.

### Evolution experiment 2

To test the effect of genetic diversity and population size (and thereby mutational supply) on adaptation to PTZ and GEN, we performed a second evolution experiment. The effect of diversity was tested by comparing the adaptation of the mixed population to the single strains H05 and H18. An increase in population size was achieved by conducting the experiment in 2 mL and 4 mL volumes while maintaining the same selective pressure (IC99.9). In total, three treatments were tested: No drug, GEN monotherapy, and PTZ monotherapy, with 20 independent replicates per treatment. Treatments were fully randomized on 48-deep-well plates. Each of the 12 strains were grown for 20 h in LB and then mixed in equal proportions (v/v). From independent mixtures, bacteria were inoculated into 48-deep-well plates containing either 2 mL or 4 mL of medium (40 µL inoculum for 2 mL and 80 µL inoculum for 4 mL). They were then grown for 24 h at 37 °C and 750 rpm, following which the plates were sonicated to break any bacterial clumps. 2% (v/v) inoculum was then transferred to another plate containing fresh medium. Growth of the bacteria was followed by measuring endpoint OD_600_ and CFU/mL over time. The experiment lasted 5 days.

### Fluctuation Assay

We performed a classical fluctuation assay to measure the rates of resistance mutations towards GEN and PTZ^36,37^. In short, one colony of the tested bacterium was inoculated into 10 ml of M9 and incubated for 20 h at 37 °C and 150 rpm. Similar to the other experiments, the genetically diverse population was prepared by mixing the overnight culture in equal ratios. Parallel cultures were initiated in 96-deep well plates, each well containing 1 mL with a 10^3^ - 10^4^ CFU/mL founding population. To control for strong bottlenecks affecting the gdPop rates, we used a larger founding population. Each strain and antibiotic combination included two independent experiments. The deep-well plates were incubated at 37 °C and 300 rpm for 20 h. Thereafter, we plated a 1:10 dilution of the cells on M9 agar plates containing 4 x MIC of either GEN or PTZ at a density of 10^5^ - 10^6^ cells/cm^2^. The mutants were counted after 48 h of growth at 37 °C. Plates containing more than 250 mutants were counted as 250 mutants. Resistance rates were determined using the Lea-Coulson model with partial plating in webSalvador 0.1^65^.

### Statistical analysis

Statistical analysis was carried out in R^66^ using packages *car*, *lawstat, lme4, vegan, MASS,* and *lmPerm*. From the first evolution experiment, the response variables ‘Time to adaptation’, ‘Robustness of adaptation’, ‘MIC on GEN’, and ‘MIC on PTZ’, were analyzed with the same statistical framework. A full factorial randomization Analysis of Variance (ANOVA) was used to assess the impact of the predictor variables ‘Meta-Treatment’, ‘Treatment’, and their interaction on the response variables. The general model of the ANOVA used was *aov(Response ~ Meta.Treatment * Treatment, data = datadf).* The model was run 10000 times with the data randomized at each run to generate a distribution of F ratios expected under the null hypothesis of no association. A *p* value was calculated as the probability of obtaining F ratios greater than or equal to the F ratios observed with the original data. Post hoc testing was carried out using pairwise Wilcoxon rank sum tests. The effect of these predictor variables on the response variable population survival was assessed using a logistic regression model defined as *glm(Survival ~ Meta.treatment + Treatment, data = survival, family = binomial)*.

The effect of the meta-treatment on Shannon diversity was analyzed using generalized linear models with the formula *lmer(Shannon ~ meta-treatment + (1|season), data = diversity).* To assess the influence of the meta-treatment and the treatment on the survival frequency of the gdPop strains, a PERMANOVA (adonis2 function of the R package *vegan*) with 10,000 permutations was conducted. To avoid modeling strain correspondence, the Bray-Curtis distance of the post-experimental survival (as a measure of beta diversity) was calculated instead of the original strain proportions, and these distances were used as the response in a full-factorial model. To assess whether pre-existing resistance and bacterial interaction significantly contribute to explaining the variance in strain survival, linear regressions were performed on the mixing and no mixing meta-treatments separately. A full factorial model was initially used, and stepwise model selection (backward elimination and forward selection) was conducted using the Akaike Information Criterion (AIC) with the stepAIC function from the R package *MASS*. Due to non-parametric and heteroscedastic residuals, the significance of the identified regression model was verified using a permutation test (lmp function from R’s *lmPerm* package), and non-significant terms were sequentially removed from the model until the final overall model and all its terms had significant *p*-values (< 0.05).

Resistance rates were compared with the likelihood ratio test, within *webSalvador* 0.1. We used a variety of initial mc, 0.1, 0.4, and 2.1 which all converged to the same likelihood ratio test statistic. All *p* values were adjusted for multiple testing with the false discovery rate.

In the second evolution experiment, the effect of Volume, Antibiotic treatment, and Strain on survival was assessed using logistic regression with the *model glm(Survival ~ Volume + Antibiotic + Strain, data=datadf, family = binomial)* while adaptation parameters were analyzed with a randomized ANOVA using the model *aov(Response ~ Volume * Strain, data = datadf*) followed by post hoc analysis using pairwise Wilcoxon rank sum tests.

## Data availability

Sequencing data have been deposited at NCBI under the BioProject number PRJNA1200785 for both Illumina HiSeq3000 short reads and the PacBio Sequel II long reads. All other data is provided in the supplementary source data files.

## Acknowledgements

We would like to thank Sara Mitri (Lausanne, Switzerland) and Kevin Foster (Oxford, UK) for their valuable advice on experimental design, and Sara Mitri, Andrew Farr (Ploen, Germany), and Alex Hall (Zuerich, Switzerland) for helpful feedback on the manuscript. We would further like to acknowledge technical help from Mandy Renner, Nadja Steinbach, Kim Flinder, and Tassja Rugenstein. We are grateful for funding from the German Research Foundation (Deutsche Forschungsgemeinschaft, DFG) within the Research and Training Group (RTG) 2501 (project 4.2 to HS and FB), within the Excellence cluster Precision Medicine in chronic Inflammation (PMI; funding under Germany’s Excellence Strategy EXC 2167-390884018, to HS and DU), individual project SCHU 1415/12 (to HS), and a Walter Benjamin grant (project BE 8013/1-1 with project number 512851323, to EBC). We also thank the Max-Planck Society for an IMPRS stipend (to AB) and a fellowship (to HS). We further thank the Max Planck-Genome-centre Cologne (http://mpgc.mpipz.mpg.de/home/) for performing all genome sequencing in this study. We also acknowledge support from the Damp Foundation (Damp Stiftung) within the SKILLED project (to HS). JB was supported by Fundação para a Ciência e Tecnologia, I.P. (FCT; CEEC contract with reference 2023.06600.CEECIND). Work in the Unterweger group was supported by the German Federal Ministry for Education and Research (grant 01KI2020).

## Author contributions

**Conceptualization:** AB, LT, EBC, HU, DU, HS

**Investigation:** AB, LT, KH, TL, JB, MH, DS, GS, FB

**Data curation:** AB, KH, TL, JB, MH

**Formal analysis:** AB, LT, JB, MH, DS, GS, FB

**Validation:** AB, KH

**Visualization:** AB, LT, JB

**Methodology:** AB, LT, KH, TL

**Supervision:** AB, LT, DU, HS

**Funding acquisition:** DU, HS

**Writing - original draft:** AB, JB, MH, FB, HS

**Writing - review and editing:** All authors

## Notes

### Competing Interest Statement

The authors have declared no competing interest.

