## Supplement for "Resistance variation and bacterial interactions shape the adaptation of a genetically diverse bacterial population to antimicrobial treatment"

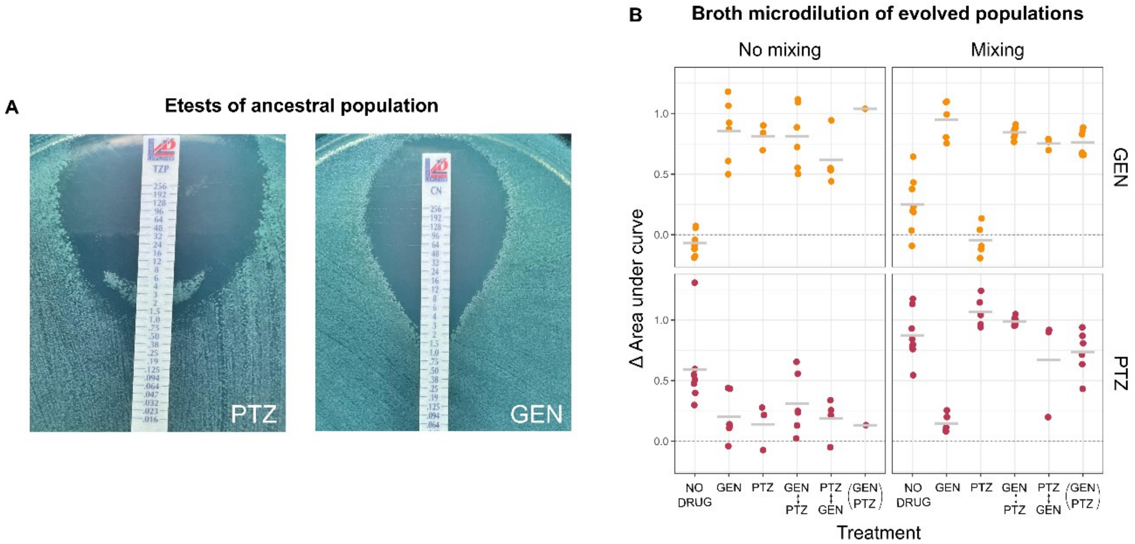

**Fig. S1: Antibiotic resistance measurements of the gdPop**. Antibiotic resistance levels of the ancestral and evolved gdPop were measured using broth microdilution and MIC test strips (Liofilchem) **(A)** The minimum inhibitory concentration (MIC) on the two antibiotics of the ancestral gdPop was determined using MIC Test strips. MIC was read at the intersection of the zone of inhibition and the test strip for GEN. For PTZ we observed a 'wing phenotype'. This is likely due to the paradoxical effect. MIC was read at the intersection of the upper edge of the wings and the strip. **(B)** Resistance of the evolved populations was also determined using broth microdilution. Frozen populations from season 14 were regrown and the MIC was measured. The number of recovered populations varied from 2-8 per treatment. Dots are the individual replicate populations while the bar is the mean. The dotted line represents the ancestral IC99.9.

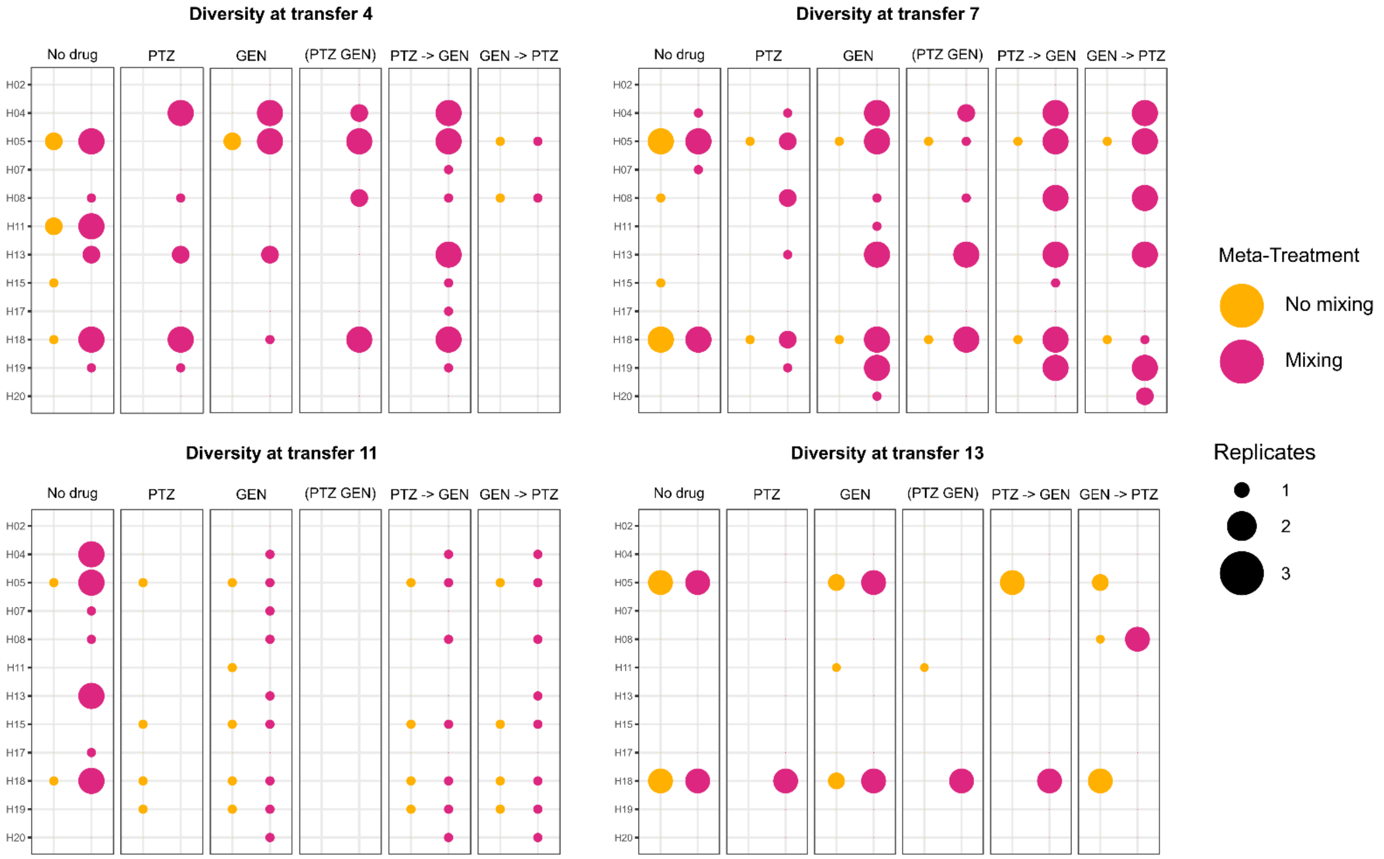

**Fig. S2: Strain diversity over time in the first evolution experiment.** Presence/absence data for the 12 strains in the gdPop was obtained through strain-specific PCRs for three replicate populations from transfers 4, 7, 11, and 13 and for every treatment and meta-treatment combination. The colour of the circles represents the meta-treatment. Size of the circles corresponds to the number of replicates the strain was observed in.

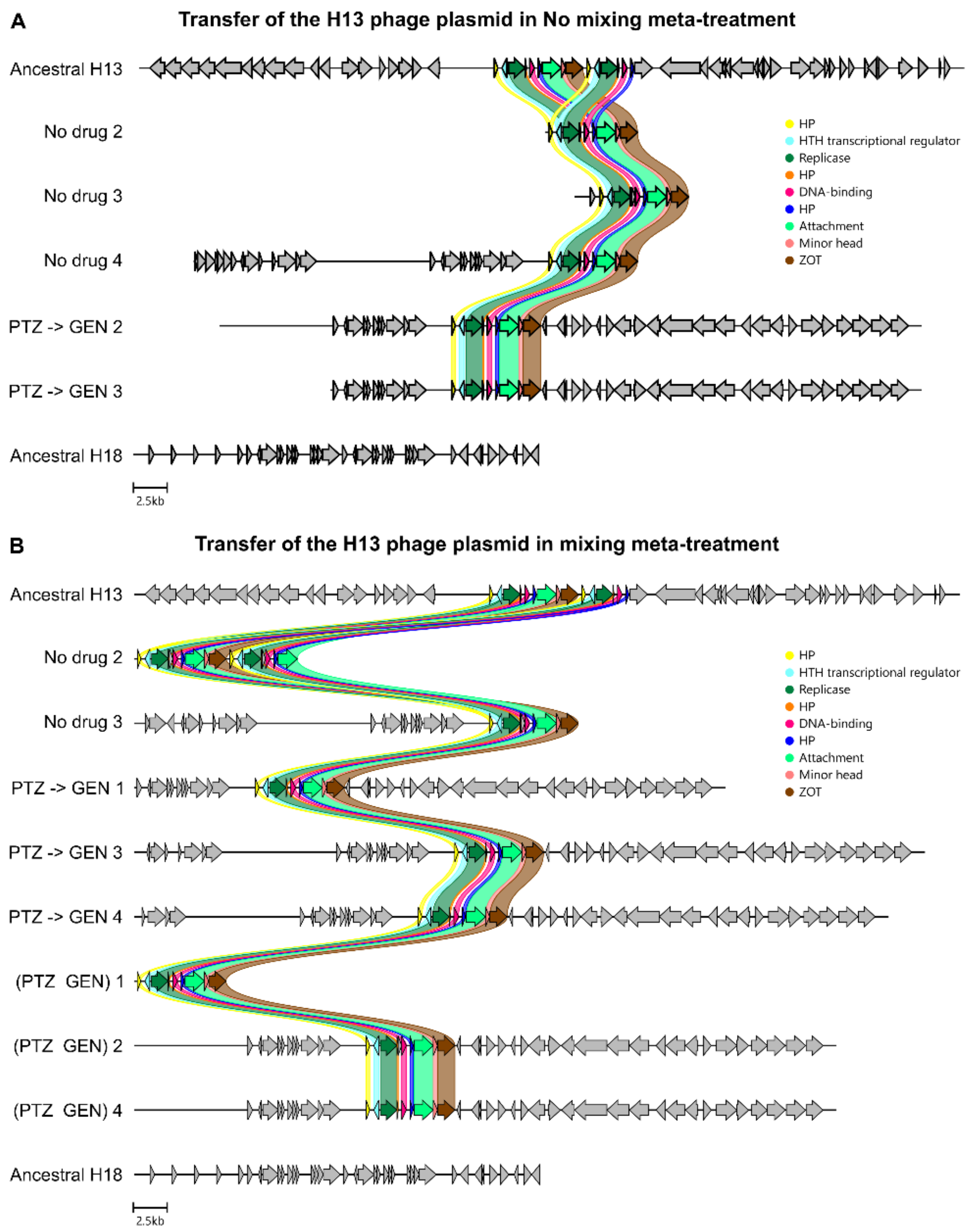

**Fig. S3: Horizontal gene transfer in the gdPop during the first evolution experiment.** A phage plasmid from ancestral H13, was observed in evolved H18 strains in several treatments in the no-mixing (A) and the mixing meta-treatments (B). The plasmid was not present in the ancestral H18. Many genes were present on the plasmid, but none was previously linked to antibiotic resistance.

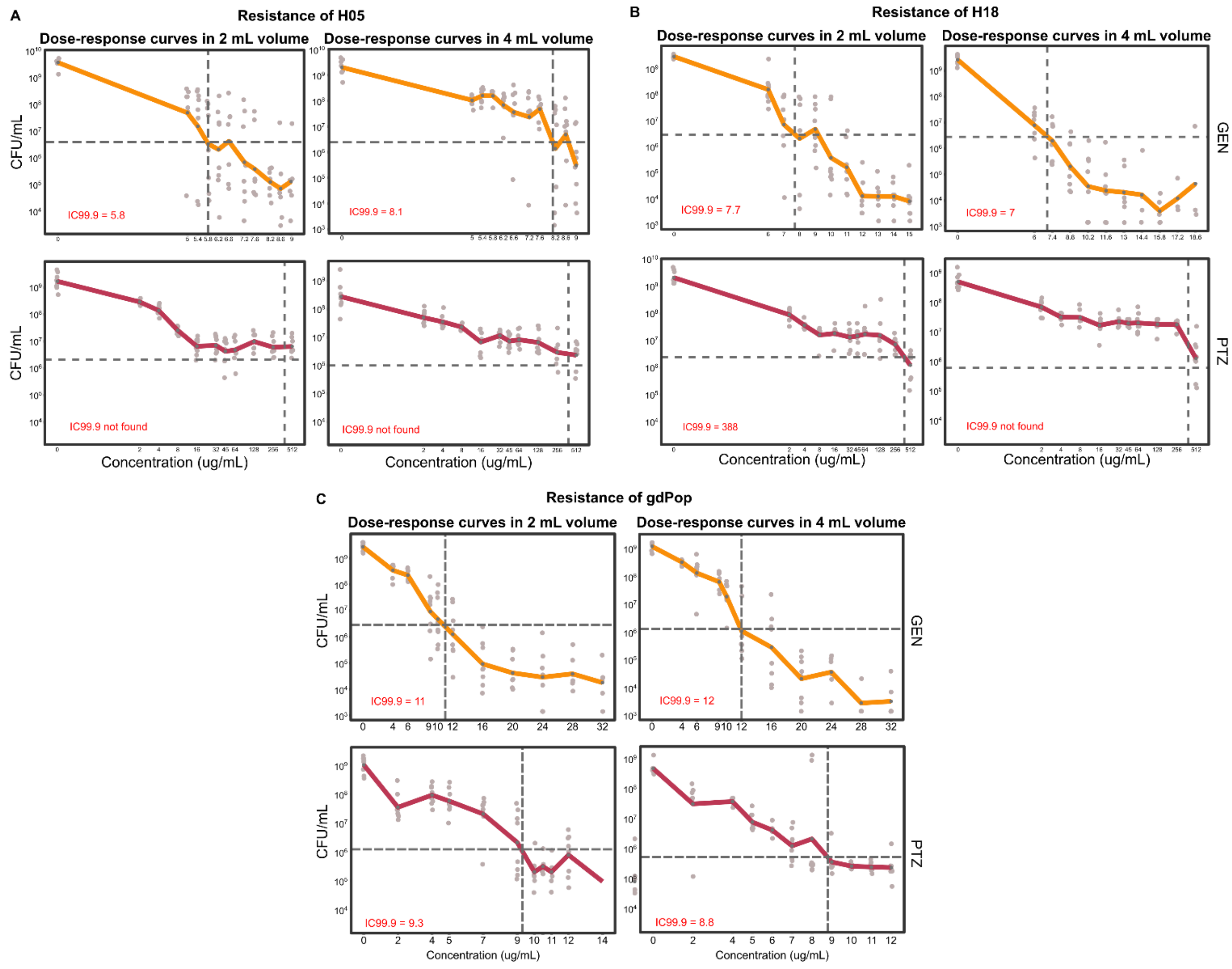

**Fig. S4: Selection concentrations of antibiotics for the second evolution experiment.** CFU-based dose response curves were performed to determine the concentration of antibiotic that killed 99.9% of the cells (IC99.9) for the single strains H05 and H18 **(A & B)** and the gdPop **(C)**. Two volumes were included for both antibiotics. Light grey dots are replicate populations and the thick coloured line is the mean. A total of 8 replicates were tested. Intersection of the dotted lines is the IC99.9. For the single strains, except for H18 in 2 mL, an IC99.9 could not be determined on PTZ. For single strains on PTZ, the concentration at which the curve plateaued (16 ug/mL) was used for the evolution experiment.

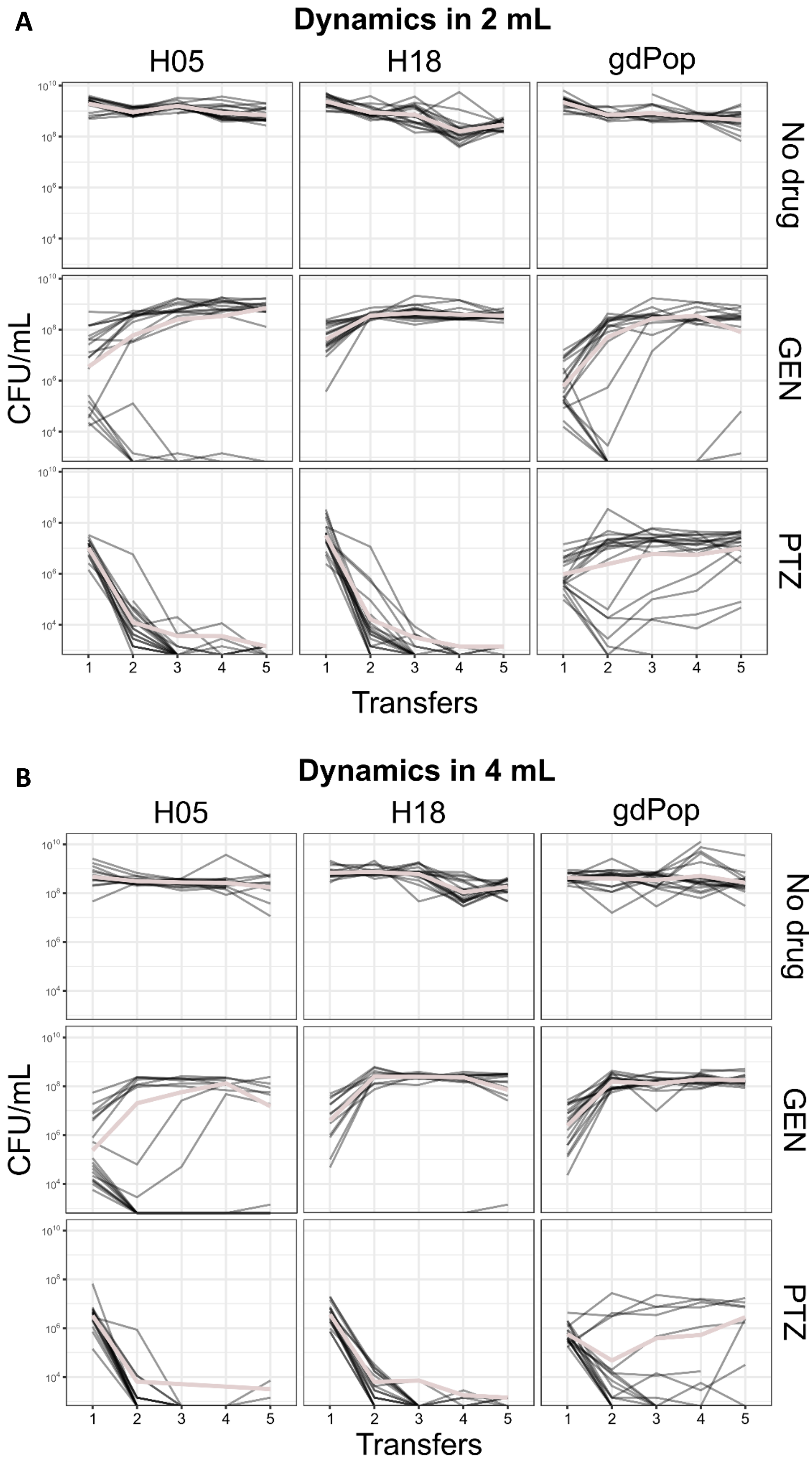

**Fig. S5: CFU over time in the second evolution experiment.** Growth dynamics under antibiotic selection for the 2 mL **(A)** and the 4 mL **(B)** volumes were followed by measuring CFU/mL at the end of each growth transfer. Thin black lines represent individual replicates, thick coloured line is the mean of all replicates. A total of 20 replicates per treatment, strain and volume combination were tested. Starting population size varied quite a bit on GEN due to the steep dose response curve of the antibiotic. Populations with a low starting population size tended to go extinct during the course of the evolution experiment.

**Supplementary Table S1. Logistic regression to assess impact of meta-treatment and treatment on survival in the first evolution experiment[a]**

| Source of variation | Estimate | *Std. Error* | *Z value* | *P* |
|---|---|---|---|---|
| (Intercept) | 18.7079 | 2404.3591 | 0.008 | 0.99379 |
| Meta-treatment (Mixing) | 2.8122 | 0.8842 | 3.181 | **0.00147** |
| Treatment (MP) | -18.9297 | 2404.3592 | -0.008 | 0.99372 |
| Treatment (SG) | -16.6876 | 2404.3593 | -0.007 | 0.99446 |
| Treatment (SP) | -18.0641 | 2404.3592 | -0.008 | 0.99401 |
| Treatment (CO) | -19.7191 | 2404.3592 | -0.008 | 0.99346 |

[a]The effects of the two factors, Meta-treatment and Treatment on the response variable survival was assessed using a logistic regression model. The analysis was performed in R version 4.3.1 using the model *glm(Survival ~ Meta-treatment + Treatment, data = survival, family = binomial)*. Significant *P* values are indicated in bold.

**Supplementary Table S2. ANOVA table for Time taken to adaptation of evolved populations[a]**

| Source of variation | *P* |
|---|---|
| Meta-treatment | **< 0.0001** |
| Treatment | **< 0.0001** |
| Meta-treatment*Treatment | **< 0.0001** |

[a]The effects of the two factors, Meta-treatment and Treatment on the response variable Time taken to adaptation was assessed using full factorial ANOVA. The analysis was performed in R version 4.3.1 using the model *aov(Time.to.adaptation ~ Meta.Treatment*Treatment, data = adaptdf)*. 10000 runs of permutation were carried out. Significant *P* values are indicated in bold.

**Supplementary Table S3. Post hoc tests comparing Treatments across Meta-treatments for Time taken to adaptation of evolved populations[a]**

| Comparison 1 | Comparison 2 | *Adjusted P* |
|---|---|---|
| No mixing Monotherapy GEN | No mixing Monotherapy PTZ | **0.0012** |
| No mixing Monotherapy GEN | No mixing (GEN -> PTZ) | **0.0132** |
| No mixing Monotherapy GEN | No mixing (PTZ -> GEN) | **0.0012** |
| No mixing Monotherapy GEN | No mixing (PTZ + GEN) | **0.0069** |
| No mixing Monotherapy GEN | Mixing Monotherapy GEN | 1 |
| No mixing Monotherapy GEN | Mixing Monotherapy PTZ | **0.0128** |
| No mixing Monotherapy GEN | Mixing (GEN -> PTZ) | **0.0012** |
| No mixing Monotherapy GEN | Mixing (PTZ -> GEN) | **0.0012** |
| No mixing Monotherapy GEN | Mixing (PTZ + GEN) | **0.0012** |
| No mixing Monotherapy PTZ | No mixing (GEN -> PTZ) | **0.0034** |
| No mixing Monotherapy PTZ | No mixing (PTZ -> GEN) | **0.0198** |
| No mixing Monotherapy PTZ | No mixing (PTZ + GEN) | 0.1917 |
| No mixing Monotherapy PTZ | Mixing Monotherapy GEN | **0.0012** |
| No mixing Monotherapy PTZ | Mixing Monotherapy PTZ | **0.0012** |
| No mixing Monotherapy PTZ | Mixing (GEN -> PTZ) | **0.0012** |

| | | |
|---|---|---|
| No mixing Monotherapy PTZ | Mixing (PTZ -> GEN) | **0.0012** |
| No mixing Monotherapy PTZ | Mixing (PTZ + GEN) | **0.0012** |
| No mixing (GEN -> PTZ) | No mixing (PTZ -> GEN) | 0.0971 |
| No mixing (GEN -> PTZ) | No mixing (PTZ + GEN) | 0.1276 |
| No mixing (GEN -> PTZ) | Mixing Monotherapy GEN | **0.0132** |
| No mixing (GEN -> PTZ) | Mixing Monotherapy PTZ | 0.1171 |
| No mixing (GEN -> PTZ) | Mixing (GEN -> PTZ) | 0.4458 |
| No mixing (GEN -> PTZ) | Mixing (PTZ -> GEN) | 0.4956 |
| No mixing (GEN -> PTZ) | Mixing (PTZ + GEN) | 0.438 |
| No mixing (PTZ -> GEN) | No mixing (PTZ + GEN) | 0.4722 |
| No mixing (PTZ -> GEN) | Mixing Monotherapy GEN | **0.0012** |
| No mixing (PTZ -> GEN) | Mixing Monotherapy PTZ | **0.0031** |
| No mixing (PTZ -> GEN) | Mixing (GEN -> PTZ) | **0.0065** |
| No mixing (PTZ -> GEN) | Mixing (PTZ -> GEN) | **0.038** |
| No mixing (PTZ -> GEN) | Mixing (PTZ + GEN) | **0.0034** |
| No mixing (PTZ + GEN) | Mixing Monotherapy GEN | **0.0069** |
| No mixing (PTZ + GEN) | Mixing Monotherapy PTZ | 0.0778 |
| No mixing (PTZ + GEN) | Mixing (GEN -> PTZ) | 0.1125 |
| No mixing (PTZ + GEN) | Mixing (PTZ -> GEN) | 0.1125 |
| No mixing (PTZ + GEN) | Mixing (PTZ + GEN) | 0.0971 |
| Mixing Monotherapy GEN | Mixing Monotherapy PTZ | **0.0128** |
| Mixing Monotherapy GEN | Mixing (GEN -> PTZ) | **0.0012** |
| Mixing Monotherapy GEN | Mixing (PTZ -> GEN) | **0.0012** |
| Mixing Monotherapy GEN | Mixing (PTZ + GEN) | **0.0012** |
| Mixing Monotherapy PTZ | Mixing (GEN -> PTZ) | 0.1122 |
| Mixing Monotherapy PTZ | Mixing (PTZ -> GEN) | **0.0034** |
| Mixing Monotherapy PTZ | Mixing (PTZ + GEN) | **0.0076** |
| Mixing (GEN -> PTZ) | Mixing (PTZ -> GEN) | **0.0221** |
| Mixing (GEN -> PTZ) | Mixing (PTZ + GEN) | 0.6532 |
| Mixing (PTZ-> GEN) | Mixing (PTZ + GEN) | **0.0079** |

[a]Post hoc comparisons following a significant permutation ANOVA were carried out using Pairwise Wilcoxon Rank Sum Tests in R version 4.3.1. *P* values were adjusted for multiple comparisons using the False Discovery Rate (FDR) method: Significant *P* values are given in bold.

**Supplementary Table S4. ANOVA table for Robustness of adaptation of evolved populations[a]**

| Source of variation | *P* |
|---|---|
| Meta-treatment | **< 0.0001** |
| Treatment | **< 0.0001** |
| Meta-treatment*Treatment | **< 0.0001** |

[a]The effects of the two factors, Meta-treatment and Treatment on the response variable Robustness of adaptation was assessed using full factorial ANOVA. The analysis was performed in R version 4.3.1 using the model *aov(Robustness.of..adaptation ~ Meta.Treatment*Treatment, data = adaptdf)*. 10000 runs of permutation were carried out. Significant *P* values are indicated in bold.

**Supplementary Table S5. Post hoc tests comparing Treatments across Meta-treatments for Robustness of adaptation of evolved populations[a]**

| **Comparison 1** | **Comparison 2** | ***Adjusted P*** |
|---|---|---|
| No mixing Monotherapy GEN | No mixing Monotherapy PTZ | **0.0014** |
| No mixing Monotherapy GEN | No mixing (GEN -> PTZ) | **0.0014** |
| No mixing Monotherapy GEN | No mixing (PTZ -> GEN) | **0.0014** |
| No mixing Monotherapy GEN | No mixing (PTZ + GEN) | **0.0014** |
| No mixing Monotherapy GEN | Mixing Monotherapy GEN | 1 |
| No mixing Monotherapy GEN | Mixing Monotherapy PTZ | **0.0014** |
| No mixing Monotherapy GEN | Mixing (GEN -> PTZ) | **0.0014** |
| No mixing Monotherapy GEN | Mixing (PTZ -> GEN) | **0.0014** |
| No mixing Monotherapy GEN | Mixing (PTZ + GEN) | **0.0014** |
| No mixing Monotherapy PTZ | No mixing (GEN -> PTZ) | **0.0114** |
| No mixing Monotherapy PTZ | No mixing (PTZ -> GEN) | 0.0579 |
| No mixing Monotherapy PTZ | No mixing (PTZ + GEN) | 0.6099 |
| No mixing Monotherapy PTZ | Mixing Monotherapy GEN | **0.0014** |
| No mixing Monotherapy PTZ | Mixing Monotherapy PTZ | **0.0014** |
| No mixing Monotherapy PTZ | Mixing (GEN -> PTZ) | **0.0014** |
| No mixing Monotherapy PTZ | Mixing (PTZ -> GEN) | **0.0014** |
| No mixing Monotherapy PTZ | Mixing (PTZ + GEN) | **0.0014** |
| No mixing (GEN -> PTZ) | No mixing (PTZ -> GEN) | 0.1008 |
| No mixing (GEN -> PTZ) | No mixing (PTZ + GEN) | **0.0081** |
| No mixing (GEN -> PTZ) | Mixing Monotherapy GEN | **0.0014** |
| No mixing (GEN -> PTZ) | Mixing Monotherapy PTZ | 1 |
| No mixing (GEN -> PTZ) | Mixing (GEN -> PTZ) | 0.7151 |
| No mixing (GEN -> PTZ) | Mixing (PTZ -> GEN) | 0.9135 |
| No mixing (GEN -> PTZ) | Mixing (PTZ + GEN) | 0.5881 |
| No mixing (PTZ -> GEN) | No mixing (PTZ + GEN) | **0.0431** |
| No mixing (PTZ -> GEN) | Mixing Monotherapy GEN | **0.0014** |
| No mixing (PTZ -> GEN) | Mixing Monotherapy PTZ | **0.0117** |
| No mixing (PTZ -> GEN) | Mixing (GEN -> PTZ) | **0.0016** |
| No mixing (PTZ -> GEN) | Mixing (PTZ -> GEN) | **0.0331** |
| No mixing (PTZ -> GEN) | Mixing (PTZ + GEN) | 0.1278 |
| No mixing (PTZ + GEN) | Mixing Monotherapy GEN | **0.0014** |
| No mixing (PTZ + GEN) | Mixing Monotherapy PTZ | **0.0014** |
| No mixing (PTZ + GEN) | Mixing (GEN -> PTZ) | **0.0014** |
| No mixing (PTZ + GEN) | Mixing (PTZ -> GEN) | **0.0014** |
| No mixing (PTZ + GEN) | Mixing (PTZ + GEN) | **0.0014** |
| Mixing Monotherapy GEN | Mixing Monotherapy PTZ | **0.0014** |
| Mixing Monotherapy GEN | Mixing (GEN -> PTZ) | **0.0014** |
| Mixing Monotherapy GEN | Mixing (PTZ -> GEN) | **0.0014** |
| Mixing Monotherapy GEN | Mixing (PTZ + GEN) | **0.0014** |
| Mixing Monotherapy PTZ | Mixing (GEN -> PTZ) | **0.0431** |
| Mixing Monotherapy PTZ | Mixing (PTZ -> GEN) | 0.5204 |
| Mixing Monotherapy PTZ | Mixing (PTZ + GEN) | **0.0431** |
| Mixing (GEN -> PTZ) | Mixing (PTZ -> GEN) | **0.0331** |
| Mixing (GEN -> PTZ) | Mixing (PTZ + GEN) | **0.0015** |

| | | |
|---|---|---|
| Mixing (PTZ-> GEN) | Mixing (PTZ + GEN) | 0.1217 |

[a]Post hoc comparisons following a significant permutation ANOVA were carried out using Pairwise Wilcoxon Rank Sum Tests in R version 4.3.1. *P* values were adjusted for multiple comparisons using the False Discovery Rate (FDR) method: Significant *P* values are given in bold.

**Supplementary Table S6. ANOVA table for resistance (MIC) on PTZ of evolved populations[a]**

| Source of variation | *P* |
|---|---|
| Meta-treatment | **< 0.0001** |
| Treatment | **< 0.0001** |
| Meta-treatment*Treatment | **< 0.0001** |

[a]The effects of the two factors, Meta-treatment and Treatment on the response variable resistance (MIC) on PTZ was assessed using full factorial ANOVA. The analysis was performed in R version 4.3.1 using the model *Anova(aov(diffANC ~ Meta.treatment * Treatment, data = PITMICdf), type = 2).* 10000 runs of permutation were carried out. Significant *P* values are indicated in bold.

**Supplementary Table S7. Post hoc tests comparing Treatments across Meta-treatments for resistance (MIC) on PTZ of evolved populations[a]**

| Comparison 1 | Comparison 2 | *Adjusted P* |
|---|---|---|
| No mixing Drug free | No mixing Monotherapy GEN | 0.957035042 |
| No mixing Drug free | No mixing Monotherapy PTZ | 0.679255159 |
| No mixing Drug free | No mixing (GEN -> PTZ) | 0.281751144 |
| No mixing Drug free | No mixing (PIT + GEN) | 0.160096357 |
| No mixing Drug free | Mixing Drug free | **0.040055443** |
| No mixing Drug free | Mixing Monotherapy GEN | 0.257929258 |
| No mixing Drug free | Mixing Monotherapy PTZ | **0.011161558** |
| No mixing Drug free | Mixing (GEN -> PTZ) | **0.011161558** |
| No mixing Drug free | Mixing (PIT + GEN) | **0.041321758** |
| No mixing Monotherapy GEN | No mixing Monotherapy PTZ | 0.597744705 |
| No mixing Monotherapy GEN | No mixing (GEN -> PTZ) | 0.112849035 |
| No mixing Monotherapy GEN | No mixing (PIT + GEN) | 0.083448072 |
| No mixing Monotherapy GEN | Mixing Drug free | **0.011161558** |
| No mixing Monotherapy GEN | Mixing Monotherapy GEN | 0.116462742 |
| No mixing Monotherapy GEN | Mixing Monotherapy PTZ | **0.011161558** |
| No mixing Monotherapy GEN | Mixing (GEN -> PTZ) | **0.011161558** |
| No mixing Monotherapy GEN | Mixing (PIT + GEN) | **0.040055443** |
| No mixing Monotherapy PTZ | No mixing (GEN -> PTZ) | 0.244674045 |
| No mixing Monotherapy PTZ | No mixing (PIT + GEN) | 0.152231068 |
| No mixing Monotherapy PTZ | Mixing Drug free | **0.032231335** |
| No mixing Monotherapy PTZ | Mixing Monotherapy GEN | 0.129322012 |
| No mixing Monotherapy PTZ | Mixing Monotherapy PTZ | **0.023772565** |
| No mixing Monotherapy PTZ | Mixing (GEN -> PTZ) | **0.014736685** |
| No mixing Monotherapy PTZ | Mixing (PIT + GEN) | 0.066422701 |

| | | |
|---|---|---|
| No mixing (GEN -> PTZ) | No mixing (PIT + GEN) | 0.330787673 |
| No mixing (GEN -> PTZ) | Mixing Drug free | 0.244674045 |
| No mixing (GEN -> PTZ) | Mixing Monotherapy GEN | 0.365631622 |
| No mixing (GEN -> PTZ) | Mixing Monotherapy PTZ | **0.016529092** |
| No mixing (GEN -> PTZ) | Mixing (GEN -> PTZ) | **0.011161558** |
| No mixing (GEN -> PTZ) | Mixing (PIT + GEN) | 0.054859563 |
| No mixing (PIT + GEN) | Mixing Drug free | 0.618716424 |
| No mixing (PIT + GEN) | Mixing Monotherapy GEN | 0.134475559 |
| No mixing (PIT + GEN) | Mixing Monotherapy PTZ | 0.134475559 |
| No mixing (PIT + GEN) | Mixing (GEN -> PTZ) | 0.152231068 |
| No mixing (PIT + GEN) | Mixing (PIT + GEN) | 0.25 |
| Mixing Drug free | Mixing Monotherapy GEN | **0.038757462** |
| Mixing Drug free | Mixing Monotherapy PTZ | **0.011161558** |
| Mixing Drug free | Mixing (GEN -> PTZ) | **0.011161558** |
| Mixing Drug free | Mixing (PIT + GEN) | **0.040055443** |
| Mixing Monotherapy GEN | Mixing Monotherapy PTZ | **0.012344417** |
| Mixing Monotherapy GEN | Mixing (GEN -> PTZ) | **0.011161558** |
| Mixing Monotherapy GEN | Mixing (PIT + GEN) | **0.041321758** |
| Mixing Monotherapy PTZ | Mixing (GEN -> PTZ) | 0.799180811 |
| Mixing Monotherapy PTZ | Mixing (PIT + GEN) | 0.108576123 |
| Mixing (GEN -> PTZ) | Mixing (PIT + GEN) | 0.181365403 |

[a]Post hoc comparisons following a significant permutation ANOVA were carried out using Pairwise Wilcoxon Rank Sum Tests in R version 4.3.1. *P* values were adjusted for multiple comparisons using the False Discovery Rate (FDR) method: Significant *P* values are given in bold.

**Supplementary Table S8. ANOVA table for resistance (MIC) on GEN of evolved populations[a]**

| Source of variation | *P* |
|---|---|
| Meta-treatment | 0.0507 |
| Treatment | **< 0.0001** |
| Meta-treatment*Treatment | **< 0.0001** |

[a]The effects of the two factors, Meta-treatment and Treatment on the response variable resistance (MIC) on PTZ was assessed using full factorial ANOVA. The analysis was performed in R version 4.3.1 using the model *Anova(aov(diffANC ~ Meta.treatment * Treatment, data = GENMICdf), type = 2).* 10000 runs of permutation were carried out. Significant *P* values are indicated in bold.

**Supplementary Table S9. Post hoc tests comparing Treatments across Meta-treatments for resistance (MIC) on GEN of evolved populations[a]**

| Comparison 1 | Comparison 2 | *Adjusted P* |
|---|---|---|
| No Mixing Drug free | No Mixing Mono GEN | **0.008730909** |
| No Mixing Drug free | No Mixing Mono PTZ | **0.011186503** |
| No Mixing Drug free | No Mixing GEN -> PTZ | **0.008730909** |
| No Mixing Drug free | No Mixing PIT + GEN | 0.068546631 |

| | | |
|---|---|---|
| No Mixing Drug free | Mixing Drug free | 0.771422597 |
| No Mixing Drug free | Mixing Mono GEN | **0.009415407** |
| No Mixing Drug free | Mixing Mono PTZ | 0.503721554 |
| No Mixing Drug free | Mixing GEN -> PTZ | **0.008730909** |
| No Mixing Drug free | Mixing PIT + GEN | **0.033793134** |
| No Mixing Mono GEN | No Mixing Mono PTZ | 1 |
| No Mixing Mono GEN | No Mixing GEN -> PTZ | 0.052879966 |
| No Mixing Mono GEN | No Mixing PIT + GEN | 0.083725322 |
| No Mixing Mono GEN | Mixing Drug free | **0.008730909** |
| No Mixing Mono GEN | Mixing Mono GEN | 0.784089919 |
| No Mixing Mono GEN | Mixing Mono PTZ | **0.008730909** |
| No Mixing Mono GEN | Mixing GEN -> PTZ | **0.008730909** |
| No Mixing Mono GEN | Mixing PIT + GEN | **0.047513125** |
| No Mixing Mono PTZ | No Mixing GEN -> PTZ | 0.231899262 |
| No Mixing Mono PTZ | No Mixing PIT + GEN | 0.220959436 |
| No Mixing Mono PTZ | Mixing Drug free | **0.011186503** |
| No Mixing Mono PTZ | Mixing Mono GEN | 0.837820524 |
| No Mixing Mono PTZ | Mixing Mono PTZ | **0.01714407** |
| No Mixing Mono PTZ | Mixing GEN -> PTZ | **0.046746144** |
| No Mixing Mono PTZ | Mixing PIT + GEN | 0.220959436 |
| No Mixing GEN -> PTZ | No Mixing PIT + GEN | 0.196536866 |
| No Mixing GEN -> PTZ | Mixing Drug free | **0.008730909** |
| No Mixing GEN -> PTZ | Mixing Mono GEN | **0.033793134** |
| No Mixing GEN -> PTZ | Mixing Mono PTZ | **0.011186503** |
| No Mixing GEN -> PTZ | Mixing GEN -> PTZ | **0.024863926** |
| No Mixing GEN -> PTZ | Mixing PIT + GEN | 0.157993872 |
| No Mixing PIT + GEN | Mixing Drug free | 0.061168654 |
| No Mixing PIT + GEN | Mixing Mono GEN | 0.077854995 |
| No Mixing PIT + GEN | Mixing Mono PTZ | 0.076387432 |
| No Mixing PIT + GEN | Mixing GEN -> PTZ | 1 |
| No Mixing PIT + GEN | Mixing PIT + GEN | 0.800234607 |
| Mixing Drug free | Mixing Mono GEN | **0.009415407** |
| Mixing Drug free | Mixing Mono PTZ | 0.691177536 |
| Mixing Drug free | Mixing GEN -> PTZ | **0.008730909** |
| Mixing Drug free | Mixing PIT + GEN | **0.032163536** |
| Mixing Mono GEN | Mixing Mono PTZ | **0.017142389** |
| Mixing Mono GEN | Mixing GEN -> PTZ | **0.008730909** |
| Mixing Mono GEN | Mixing PIT + GEN | **0.040500536** |
| Mixing Mono PTZ | Mixing GEN -> PTZ | **0.008730909** |
| Mixing Mono PTZ | Mixing PIT + GEN | **0.040500536** |
| Mixing GEN -> PTZ | Mixing PIT + GEN | 0.46170086 |

[a]Post hoc comparisons following a significant permutation ANOVA were carried out using Pairwise Wilcoxon Rank Sum Tests in R version 4.3.1. *P* values were adjusted for multiple comparisons using the False Discovery Rate (FDR) method: Significant *P* values are given in bold.

**Supplementary Table S10. Strain-specific primers to detect individual strains of the mixed population**[1]

| Primer | Name | Sequence |
|---|---|---|
| H02_forward | H02_P1_F | GGA CGG TTG CCA CAA TGT TC |
| H02_reverse | H02_P1_R | CTT GCC GTT GAT AGC GAT GC |
| H04_forward | H04_P1_F | CGT TCA GCG CTA CTT CAT CG |
| H04_reverse | H04_P1_R | CTC CTG CTC TGG TTT CCT GG |
| H05_forward | H05_P1_F | CGC TGC TAC TCT CCA TCC TG |
| H05_reverse | H05_P1_R | CTG AGG CCT GGC TGA TGT AG |
| H07_forward | H07_P1_F | ACT GAA CGT AGC CAT GGA GC |
| H07_reverse | H07_P1_R | CGC CAG GTT TAG AGC CAG AG |
| H08_forward | H08_P1_F | CTG ATG CCG GAG ATC ACC TG |
| H08_reverse | H08_P1_R | GTG AGC GGT TCC AGG GAT G |
| H11_forward | H11_P1_F | GGC CGT ATC CTT TAC CTC CG |
| H11_reverse | H11_P1_R | ATC AGC AGT TCG TCA GCC TC |
| H13_forward | H13_P1_F | ACG GCA GTC TCC ATG AGT TC |
| H13_reverse | H13_P1_R | GCT CCG ATA CCG TAG AAG CC |
| H15_forward | H15_P1_F | AGA TGT TCC AGA AGC ZGC GC |
| H15_reverse | H15_P1_R | GAG AGT TTG CGT CTA TGT GC |
| H17_forward | H17_P1_F | GAG CGA CGA ATT CCT GCA AC |
| H17_reverse | H17_P1_R | GAC CAT CTC CAC TAG CTC GC |
| H18_forward | H18_P1_F | CCG CAG CCT ATC TCT ACA CG |
| H18_reverse | H18_P1_R | CCG GGT AGA GTT GGA TCA CG |
| H19_forward | H19_P1_F | TTG ACG ATC CTC AGG TGC TG |
| H19_reverse | H19_P1_R | CGA GGT GAT CTG CGA CTA CC |
| H20_forward | H20_P1_F | CCG GAC AGT TAC ACC TCG TC |
| H20_reverse | H20_P1_R | TCG GCG TCA TGT GTC AGT AC |

[1] The name indicates the strain, for which the primer pair is specific.

**Supplementary Table S11. Generalized linear model (mixed effects) to assess the effect of Meta-treatment and Treatment on Shannon diversity over time in the first evolution experiment[a]**

| **Source of variation** | | | | | |
|---|---|---|---|---|---|
| Random effects | **Variance** | | | | |
| Season (intercept) | 0.2785 | | | | |
| Residual | 0.1741 | | | | |
| Fixed effects | **Estimate** | ***Std. Error*** | ***df*** | ***t statistic*** | ***P*** |
| (Intercept) | 0.859 | 0.284 | 3.806 | 3.020 | 0.041 |
| Meta-treatment (Mixing) | 0.385 | 0.096 | 78.114 | 4.009 | **0.0001** |
| Treatment (GEN only) | 0.026 | 0.136 | 78.057 | 0.192 | 0.848 |
| Treatment (PTZ only) | -0.212 | 0.162 | 78.043 | -1.311 | 0.193 |
| Treatment (PTZ -> GEN) | 0.025 | 0.141 | 78.043 | 0.182 | 0.856 |
| Treatment (GEN -> PTZ) | 0.002 | 0.144 | 78.054 | 0.019 | 0.984 |
| Treatment (PTZ + GEN) | -0.277 | 0.158 | 78.182 | -1.743 | 0.085 |

[a]The effects of the two factors, Meta-treatment and Treatment on the response variable Shannon diversity over time was assessed using a generalized linear mixed effects model. The analysis was performed in R version 4.3.1 using the model *lmer(shan ~ mixing + treatment + (1|season), data = diversity)* and the package lmerTest. Significant *P* values are indicated in bold.

**Supplementary Table S12. PERMANOVA to assess the effect of Meta-treatment and Treatment on Strain abundance at transfer 14 in the first evolution experiment[a]**

| **Source of variation** | ***df*** | ***Sum of sq*** | ***F statistic*** | ***P*** |
|---|---|---|---|---|
| Meta-treatment | 1 | 0.317 | 3.6889 | **0.026** |
| Treatment | 5 | 6.937 | 16.1135 | **<0.0001** |
| Interaction | 5 | 8.822 | 20.4910 | **<0.0001** |
| Residual | *62* | *5.339* | | |
| Total | *73* | *21.417* | | |

[a]To assess whether combinations of meta-treatment and treatment influence the survival of individual strains, a PERMANOVA was performed using the adonis2 function from the vegan package in R. Beta diversities calculated as Bray-Curtis distances based on strain abundances have been used as response variable.

**Supplementary Table S13. Permutation linear regression (meta-treatment = mixing) to assess impact of resistance (MIC) and strain interactions (Focal -> Others and Others -> Focal) on strain abundance at transfer 14 of evolution experiment 1[a]**

| Coefficient | *Estimate* | *P* |
|---|---|---|
| MIC(PTZ)*Others -> Focal | 0.004 | **0.045** |
| Focal -> Others | **-0.002** | **< 0.0001** |

[a]The abundance of each strain at the end of the experiment can depend on various physiological factors, such as its resistance to the antibiotics GEN and PTZ, increased or decreased growth due to secretions from other strains (Others -> Focal), the effect of its own secretions on other strains (Focal -> Others), its growth rate (AUC mean), and all potential interactions among these factors. Due to the lack of parametric test assumptions, a permuted linear regression was used to determine which predictors (including their interactions) most significantly explained the mean abundance of each strain at the end of the experiments. Predictors were iteratively removed from the model in a stepwise fashion, starting with the least significant, until only significant parameters remained. These parameters and their significance levels are presented in the table for the meta-treatment “mixing”.

**Supplementary Table S14. Permutation linear regression (meta-treatment = no mixing) to assess impact of resistance (MIC) and strain interactions (Focal -> Others and Others -> Focal) on strain abundance at transfer 14 of evolution experiment 1[a]**

| Coefficient | *Estimate* | *P* |
|---|---|---|
| MIC(PTZ) | 0.026 | **<0.0001** |
| MIC(PTZ)*Others -> Focal | **-0.001** | **<0.0001** |
| Focal -> Others | **-0.002** | **<0.0001** |
| MIC(PTZ)*Others -> Focal* Focal -> Others | **<0.001** | **<0.0001** |

A permutation linear regression was performed as described above. These parameters and their significance levels are presented in the table for the meta-treatment “no mixing”.

**Supplementary Table S15. Likelihood Ratio tests comparing PTZ and GEN resistance rates for H05, H18 and gdPop[a]**

| Strain | Comparison | LRT statistic | *Adjusted P* |
|---|---|---|---|
| H05 | PTZ - GEN | 924.4955 | **<0.0001** |
| H18 | PTZ - GEN | 358.9571 | **<0.0001** |
| gdPop | PTZ - GEN | 106.0776 | **<0.0001** |

[a] Resistance rates were compared with the likelihood ratio test, within webSalvador 0.1 (ref). We used a variety of initial mc, 0.1, 0.4, and 2.1 which all converged to the same likelihood ratio test statistic. All *p* values were adjusted for multiple testing with the *false discovery* rate.

**Supplementary Table S16. Logistic regression to assess impact of Volume, Diversity, and Antibiotic treatment on survival in the validation evolution experiment[a]**

| Source of variation | Estimate | *Std. Error* | *Z value* | *P* |
|---|---|---|---|---|
| (Intercept) | 1.012 | 0.310 | 3.261 | **0.001** |
| Volume (4 mL) | -0.7756 | 0.3749 | -2.069 | **0.0385** |
| Antibiotic (PTZ) | -3.7479 | 0.4779 | -7.843 | **<0.0001** |
| Strain (gdPop) | 2.1280 | 0.482 | 4.408 | **<0.0001** |

[a]The effects of the three factors, Volume, Diversity, and Antibiotic treatment on the response variable survival was assessed using a logistic regression model. The analysis was performed in R version 4.3.1 using the model *glm(adapted ~ Volume + Antibiotic + Strain, data = wholedata_sinco, family = binomial)*. Since we were interested in testing the effect of single strain vs gdPop (to assess diversity) H05 and H18 data were combined for the purposes of the analysis. Significant *P* values are indicated in bold.

**Supplementary Table S17. ANOVA table for Time taken to adaptation of populations evolved to GEN in the validation evolution experiment[a]**

| Source of variation | *P* |
|---|---|
| Volume | 0.5911 |
| Strain | 0.0849 |
| Volume*Strain | 0.0565 |

[a]The effects of the two factors, Volume and Strain on the response variable Time taken to adaptation was assessed using full factorial ANOVA. The analysis was performed in R version 4.3.1 using the model *Anova(aov(Time.to.adaptation ~ Volume * Strain, data = adaptdf), type =2)*. 10000 runs of permutation were carried out. Significant *P* values are indicated in bold.

**Supplementary Table S18. ANOVA table for Robustness of adaptation of populations evolved to GEN in the validation evolution experiment[a]**

| Source of variation | *P* |
|---|---|
| Volume | **0.0078** |
| Strain | **0.0187** |
| Volume*Strain | 0.1778 |

[a]The effects of the two factors, Volume and Strain on the response variable Robustness of adaptation was assessed using full factorial ANOVA. The analysis was performed in R version 4.3.1 using the model *Anova(aov(Robustness.of.adaptation ~ Volume * Strain, data = adaptdf), type =2)*. 10000 runs of permutation were carried out. Significant *P* values are indicated in bold.

**Supplementary Table S19. Post hoc tests comparing different strains and gdPop for Robustness of adaptation of populations evolved to GEN[a]**

| **Strain 1** | **Strain 2** | ***P*** |
|---|---|---|
| H05 | H18 | 0.107 |
| H05 | gdPop | **0.019** |
| gdPop | H18 | 0.107 |

[a]Post hoc comparisons following a significant permutation ANOVA were carried out using Pairwise Wilcoxon Rank Sum Tests in R version 4.3.1. *P* values were adjusted for multiple comparisons using the False Discovery Rate (FDR) method: Significant *P* values are given in bold.
